# Phenotype control techniques for Boolean gene regulatory networks

**DOI:** 10.1101/2023.04.17.537158

**Authors:** Daniel Plaugher, David Murrugarra

## Abstract

Modeling cell signal transduction pathways via Boolean networks (BNs) has become an established method for analyzing intracellular communications over the last few decades. What’s more, BNs provide a course-grained approach, not only to understanding molecular communications, but also for targeting pathway components that alter the long-term outcomes of the system. This has come to be known as *phenotype control theory*. In this review we study the interplay of various approaches for controlling gene regulatory networks such as: algebraic methods, control kernel, feedback vertex set, and stable motifs. The study will also include comparative discussion between the methods, using an established cancer model of T-Cell Large Granular Lymphocyte (T-LGL) Leukemia. Further, we explore possible options for making the control search more efficient using reduction and modularity. Finally, we will include challenges presented such as the complexity and the availability of software for implementing each of these control techniques.

## 1 Introduction and Motivation

In biology, phenotypes represent observable features such as apoptosis, proliferation, senescence, autophagy, and more. Mathematically, a *phenotype* is associated with a group of attractors where a subset of the system’s variables have a shared state. We define an *attractor* as a set of states from which there is no escape as the system evolves, and an attractor with a singleton state is called a *fixed point*. These shared states are then used as biomarkers that indicate diverse hallmarks of the system that one might view as rolling a ball down Waddington’s epigenetic landscape [1]. Thus, *phenotype control* is the ability to drive the system to a predetermined phenotype from any initial state by inducing the appropriate gene knockouts or knock-ins [2].

One way mathematicians are able to assist biological researchers is through modeling cell signal transduction pathways. However, these pathways can be highly complex due to signaling motifs like feedback loops, crosstalk, and high-dimensional nonlinearity [3]. To address these complexities, mathematical modelers have developed many strategies for creating and analyzing networks, traditionally classified based on the time and population of gene products. For instance, there are techniques for continuous population with continuous time such as ordinary differential equations [3, 4], discrete population with continuous time such as the Gillespie formulation [5, 6], and discrete population with discrete time such as BNs, logical models, and also their related stochastic counterparts [7–11]. There are also numerous well developed statistical, agent based, and PDE models which are outside the scope of this review [2]. For this review, the framework of choice utilizes Boolean networks.

Today increasingly extensive effort is dedicated to understanding more than just the cancer cells themselves. Modelers have developed multicellular models including cancer, stromal, immune, and other cells to study the interplay between cancer cells and their surrounding tumor microenvironment [12–15]. These models are typically referred to as *multiscale* because they integrate interactions at differing size and time scales, making it possible to simulate clinically relevant spatiotemporal scales, and at the same time simulate the effect of molecular drugs on tumor progression [16–21]. The high complexity of these models generates challenges for model validation such as the need to estimate too many model parameters and controlling variables at differing scales [12, 22].

Understanding such mechanisms is quite convoluted and is not presently well-established. Even though multiscale or hybrid models would likely provide more realistic simulations, there are currently no control methods that apply directly to such models [2, 12, 22]. For this reason, we elect to utilize Boolean networks because they provide a course-grained description of gene regulatory networks without the need for tedious parameter discovery [23]. This framework would also allow for approximating multistate, multiscale, or even continuous systems by projecting into a Boolean setting for analysis [12, 24, 25]. While there are many techniques available for controlling Boolean networks, we will highlight methods that provide overarching theory, as well as some emerging techniques. These methods include computational algebra [26, 27], control kernel [28,29], feedback vertex set [30,31], and stable motifs [32], where each tactic provides a complimentary approach depending on the information available [22,33]. We will also include techniques to address efficiency with network modularity [34] and reduction [33, 35–37].

Phenotype control has two main distinguishing features. Its objectives are related to dynamical attractors of highly nonlinear systems, and it focuses on open-loop interventions. These types of interventions are instances where the protocol is not adjusted based on the state of the system, inducing the control only at the front end. This is contrasted with optimal control, where the goal is to find a control policy that specifies the ideal control action for each state [38–42]. Thus, phenotype control theory is primarily concerned with identifying key markers of the system that aid in understanding the various functions of cells and their molecular mechanisms.

The format of this review will be as follows: Section 2 will provide an initial overview of the methods with discussion of overlapping features and application to a known cancer model (Section 2.1), Section 3 will lay out the different techniques used to find target controls, Section 4 will discuss methods to make the target discovery problem more efficient, Section 5 will address limitations and open problems, Section 6 will have some concluding thoughts and discussion. Finally, readers can find helpful information in the Appendix including: toy models for basic examples of each method (Section 7.1), foundational principles for finite dynamical systems (Section 7.2), simulation techniques of suggested targets (Section 7.3), software with tutorials and how-to documentation (Section 7.4), and lastly Section 7.5 has supplementary tables.

## 2 Overview of Control Methods

Depending on the specific aims and information available, Table 1 provides a set of complementary approaches for phenotype control and their key features. For instance, if you only have access to the wiring diagram, then feedback vertex set (FVS) is an option for global stabilization. If you have the Boolean rules, and if the objective is to drive the system into one of the existing attractors, then stable motifs (SM) are an option. If you have the Boolean rules, and if the objective is to create a new attractor or to block existing attractors, then algebraic methods (AM) (also called computational algebra -CA) are an option.

**Table 1:** Phenotype methods and their features. This table contains a summary of the target identification techniques discussed, as well as their key features. Namely, we summarize their objectives, induced control actions, and the necessary components to use each method. Software for these methods can be found in the Appendix.

| Method | Control Objective(s) | Control Action(s) | Requirements | Refs |
| --- | --- | --- | --- | --- |
| Algebraic Methods | Transform transient state into a steady state; Transform steady state into a transient state; Eliminate transition between two states | Assign node to specified value; Activate or inhibit specific edge | Regulatory network structure; Boolean functions written as polynomials | [26, 43] |
| Control Kernel | Force the system to have one stable attractor | Assign node to specified value | Boolean functions written as polynomials | [28, 29] |
| Feedback Vertex Set | Force the system to have one stable attractor | Assign node to its value in the target attractor | Regulatory network structure; Node activities in target attractor | [30, 44] |
| Stable Motifs | Force any initial state toward a pre-existing attractor; Transform a steady-state into a transient state | Assign node to its stable motif value; Inhibit interaction to disrupt stable motif of a steady-state | Regulatory network structure; Boolean rules written in DNF | [32] |

Despite the shared goals of these methods, each seeks distinct control objectives. They are each based on specific mathematical structures and lack a common theoretical framework that allows their complementary and synergistic application. Yet, we clearly see overlapping outcomes between methods. For example, it has been shown that the FVS establishes the upperbound for the magnitude of targets required to control the system [29]. Indeed, we observe that, among methods using pre-existing attractors, the control sets for CA and SM are subsets of the larger FVS results. On the other hand, CA and CK appear to produce minimal sets. Further, the CA and SM methods can produce the same results, or CA can be a subset of SM. See Tables 2 and 3.

**Table 2:**
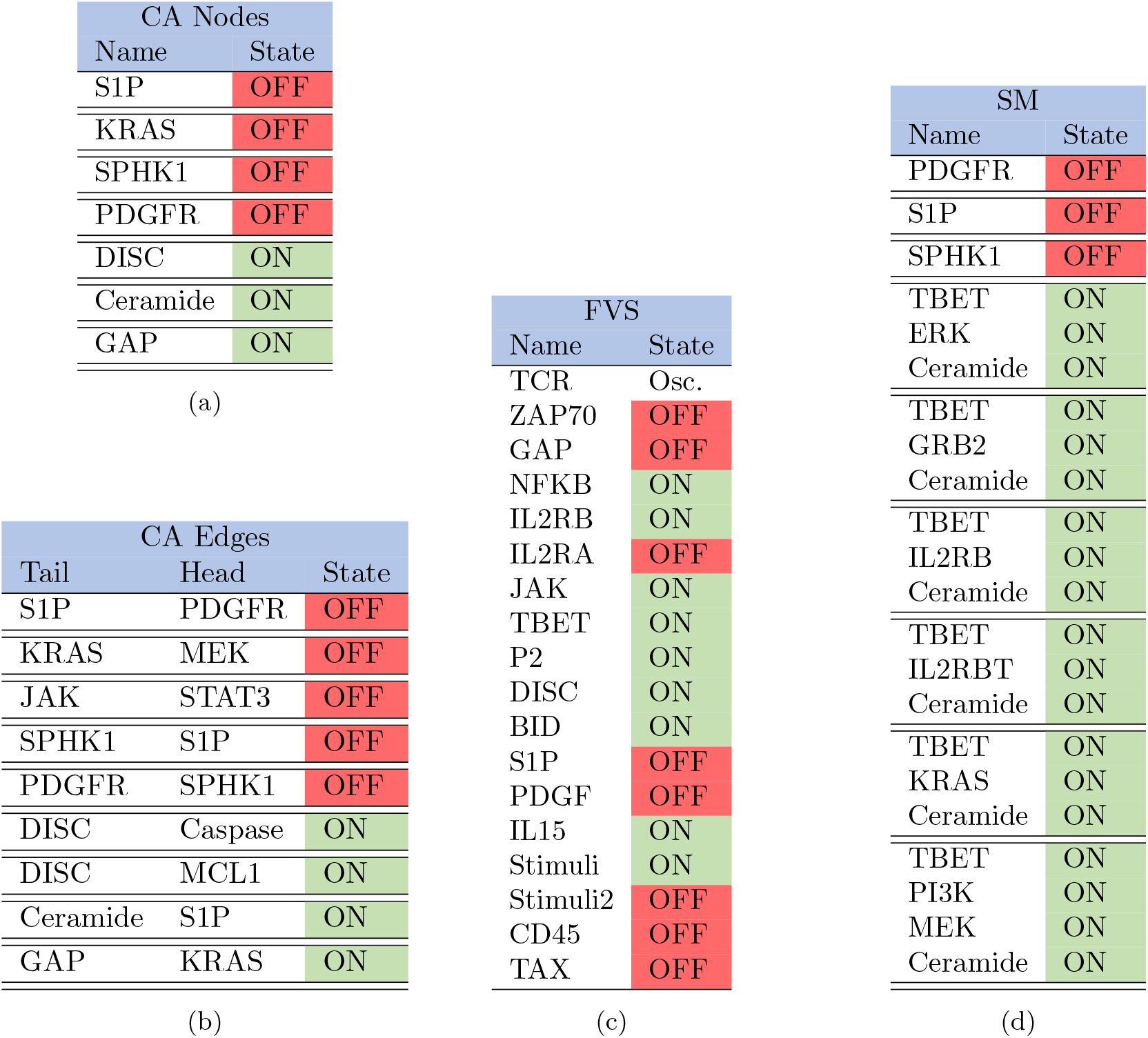
Large T-LGL target tables. Here we list the control targets for the larger T-LGL model, where control sets are separated by double horizontal bars such that Table 2a contains seven singleton controls, Table 2b contains nine singleton controls, Table 2c contains one set of 18 controls (some of which are unnecessary), and Table 2d contains three singleton controls, five triple control sets, and one quadruple control set [2].

**Table 3:** Reduced T-LGL target tables. As before, we list the control targets for the small T-LGL model, where control sets are separated by double horizontal bars such that Table 3a contains two singleton controls, Table 3b contains three singleton controls, Table 3c contains one singleton, Table 3d contains four sets of dual controls, and Table 3e contains two singleton controls [2].

| CA Edges |  |  |
| --- | --- | --- |
| Tail | Head | State |
| Ceramide | DISC | ON |
| Ceramide | S1P | ON |

(a)
| CA Nodes |  |
| --- | --- |
| Name | State |
| S1P | OFF |
| DISC | ON |
| Ceramide | ON |

(b)
| CK |  |
| --- | --- |
| Name | State |
| S1P | OFF |

(c)
| FVS |  |
| --- | --- |
| Name | State |
| Ceramide | ON |
| DISC | ON |
| Ceramide | ON |
| FLIP | OFF |
| S1P | OFF |
| DISC | ON |
| S1P | OFF |
| FLIP | OFF |

(d)
| SM |  |
| --- | --- |
| Name | State |
| S1P | OFF |
| Ceramide | ON |

However, a key unique feature of CA is the creation of new attractors, while other methods discussed rely on pre-existing attractors. This then leads to the potential for new target discovery as the long-term objectives change. Further, CA sets out to solve a system of polynomial equations, whereas FVS and SM rely on strongly connected components to find their targets. To explicitly see these connections, consider the following example.

### 2.1 Case Study: T-Cell Large Granular Lymphocyte (T-LGL) Leukemia

T-cell large granular lymphocyte (T-LGL) leukemia is a blood cancer in which there is an anomalous surge in white blood cells, called T-cells. Cytotoxic T-cells are part of the immune system that fight against antigens, even by killing cancer cells. These T-cells release specific cytokines that alter how the immune system responds to external agents by way of recruiting particular immune cells to fight infection, promoting antibody production, or inhibiting the activation and proliferation of other cells [32]. Once their job is complete they undergo controlled cell-death, however, T-LGL leukemia occurs when these T-cells evade apoptosis and maintain proliferation [2]. There are currently no standards of treatment established, however options include immunosuppressive therapy (such as methotrexate), oral cyclophosphamide (an alkylating agent), or cyclosporine (an immunomodulatory drug) [45]. Since there continues to be a search for standard therapies for this disease, the identification of potential therapeutic targets is essential.

In [46], a Boolean dynamic model was constructed consisting of a network of sixty nodes indicating the cellular location, molecular components, and conceptual nodes. For the sake of our analysis, we use the Boolean rules in Table 8 (see Appendix). The main inputs to the network are “Stimuli”, which represent virus or antigen stimulation, and the main output node is “Apoptosis”. Model analysis revealed that the system contains three attractors, two of which are diseased and the other is healthy (determined by apoptosis activation). Table 2 lists the control targets discovered by each of the respective methods for the large T-LGL model, with the objective of activating apoptosis. Individual control methods are found in Tables (2a) - (2d), and control sets are separated by double horizontal bars. Note that the CK method did not produce results for the large model because of its size [2].

Likewise, an analysis of a smaller (reduced) model of T-LGL can also be useful [11, 46]. Model analysis indicated that the reduced model in Figure 1 contains two fixed points, one healthy and one diseased. Regulatory functions for the small T-LGL model can be found in Appendix Table 7. Tables (3a) - (3e) list the control targets discovered by each of the respective methods for the small T-LGL model, with the objective of activating apoptosis. The control sets are separated by double horizontal bars as before [2].

**Figure 1:**
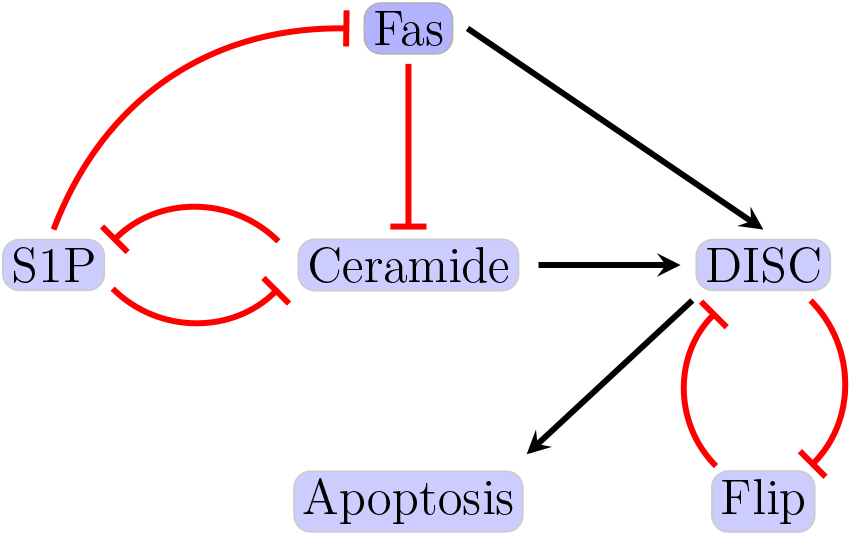
Reduced T-LGL network. The figure shown here indicates the smaller (reduced) T-LGL model, where black barbed arrows indicate signal expression and while red bar arrows indicate suppression [2].

**Figure 2:**
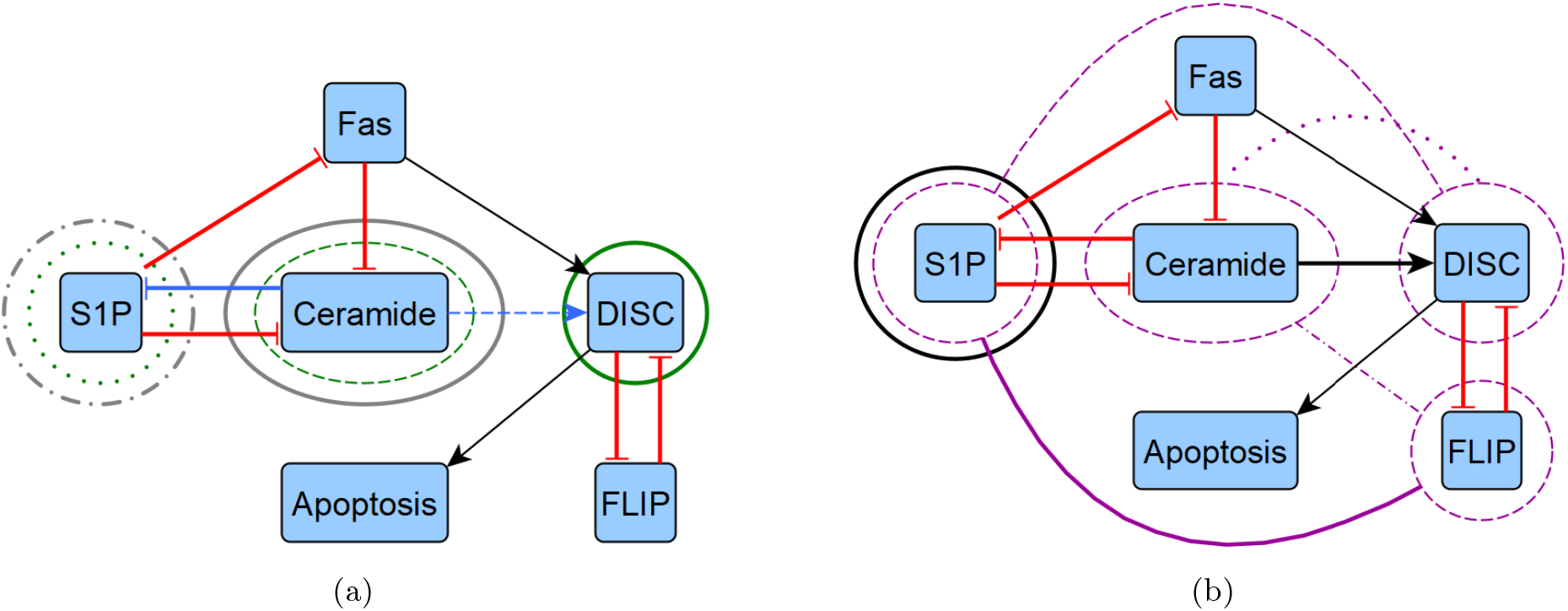
Reduced T-LGL network target overlaps. We highlight the overlapping control targets from Table 3 by overlaying them with the reduced T-LGL wiring diagram from Figure 1, shown in two diagrams to avoid excessive noise. (a) We show instances of CA edge (blue), CA node (green), and SM (grey). (b) We show instances of CK (black) and FVS (purple). Note that FVS has combinatorial controls with connecting arches, where others are strictly singleton.

For both large and reduced models, we see that FVS provides an upper bound for the amount of targets needed to achieve network control, whereas CA and CK can provide minimal sets.

## 3 Description of Control Methods

### 3.1 Algebraic Methods (CA)

The method based on computational algebra described in [26, 43] seeks two types of controls: nodes and edges. These can be achieved biologically by blocking effects of the products of genes associated with nodes, or by targeting specific gene communications (see Figure 3). The identification of control targets is achieved by encoding the nodes (or edges) of interest as control variables within the functions. Then, the control objective is expressed as a system of polynomial equations that is solved by computational algebra techniques. Though node and edge control are similar, they provide a range of biological options. One reason is that node control requires an entire node to be knocked out (or knocked-in), but edge control simply requires an edge communication to be blocked (or continually expressed) [2].

**Figure 3:**
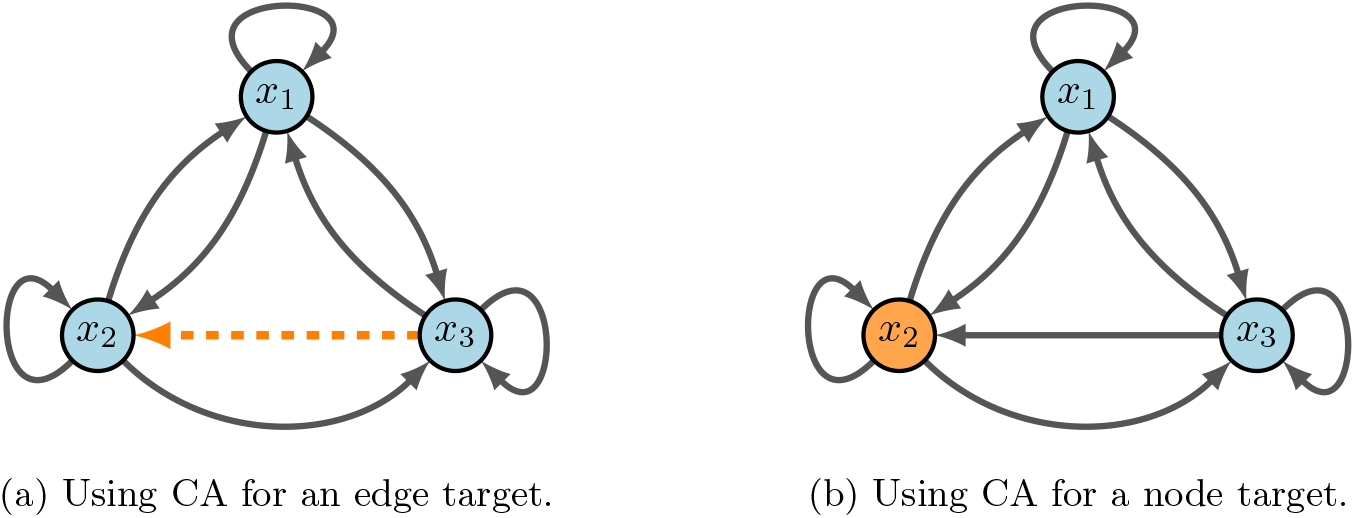
CA diagram. Here, we show a toy model that emphasizes the difference between node and edge control. The key difference with edge control (b), is that all other communcations are maintained. Whereas, node control removes every signal associated with the given target.

Let the function 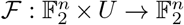 denote a Boolean network with control, where *U* is a set of all possible controls. Then, for *u* ∈ *U*, the new system dynamics are given by *x*(*t* + 1) = ℱ(*x*(*t*), *u*). That is, each coordinate *u*_*i,j*_ ∈ *u* encodes the control of edges as follows: consider the edge *x*_*i*_ → *x*_*j*_ in a given wiring diagram. Then, we can encode this edge as a control edge by the following function:

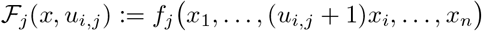

which gives

- Inactive control:

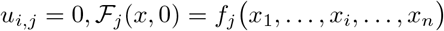
- Active control (edge deletion):

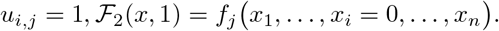

The definition of edge control can therefore be applied to many edges, obtaining 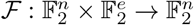 where is the number of edges in the diagram. Next, we consider control of node *x*_*i*_ from a given diagram. We can encode the control of node *x*_*i*_ by the following function:

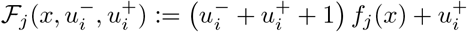

which yields

- Inactive control:

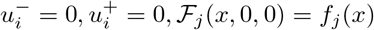
- Node *x*_*i*_ deletion:

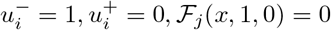
- Node *x*_*i*_ expression:

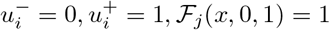
- Negated function value (irrelevant for control):

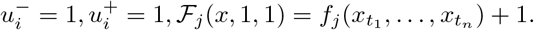

Using these definitions, we can achieve three types of objectives. Let *F* = (*f*_1_, …, *f*_*n*_) : 𝔽^*n*^ → *𝔽*^*n*^ where 𝔽 = {0, 1} and *μ* = {*μ*_1_, …, *μ*_*n*_} is a set of controls. Then we may:

- *Generate new attractors*. If *y* is a desirable state (i.e. apoptosis), but it is not currently an attractor, we find a set *μ* so that we can solve

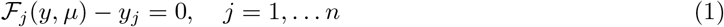
- *Block transitions or remove attractors*. If *y* is an undesirable attractor (i.e. proliferation), we want to find a set *μ* so that ℱ(*y, μ*) ≠ *y*. In general, we can use this framework to avoid transitions between states (say *y* → *z*) so that ℱ(*y, μ*) ≠ *z*. So we can solve

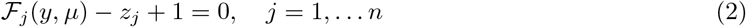
- *Block regions*. If a particular value of a variable, say *x*_*k*_ = *a*, triggers an undesirable pathway, then we need all attractors to satisfy *x*_*k*_ ≠ *a*. So we find a set *μ* so that the following system has no solution

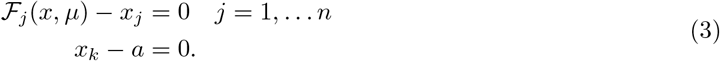

A subtle change in notation requires attention, because we have now used *x* to indicate variables rather than specific values.

Notably, the Boolean functions *F* must be written as polynomials. To complete the control search we then compute the Gr*ö*bner basis of the ideal associated with the given objective. For example, if we generate new attractors, we find the Gr*ö*bner basis for the ideal

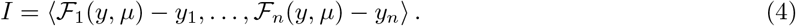

Therefore, we can determine all controls that solve the system of equations and detect combinatorial actions for the given model [2].

### 3.2 Control Kernel (CK)

A *control kernel (CK)* is defined as a set of nodes of minimal order whose pinning reshapes the dynamics such that the basin of attraction of attractor *A* becomes the entire configuration space. There are three main contributors to the CK: input nodes (nodes with identity function as the updating rule), distinguishing nodes (subset of nodes where a pinning exists that is both compatible with attractor *A* and incompatible with the other initial attractors of the network), and additional nodes (minimal distinguishing node sets that are needed to remove additional attractors). Note that input and distinguishing nodes provide only a lower bound to CK size because the pinning procedure can create new attractors [2, 29].

To compute CKs, first start with pinning input nodes. Then a brute-force method is used to loop over sets of distinguishing nodes of increasing size for each attractor. A CK has been found when no other attractors exist after pinning. Uncontrollable complex attractors are identified by pinning all constant nodes. If more than one attractor remains, then the cycle does not have a CK [29]. CK discovery works well for small networks, however, larger networks prove more difficult due to the brute-force nature of the algorithm. In fact, the scaling of the set cardinality is logarithmic based on the number of attractors in the network [2, 29].

### 3.3 Feedback Vertex Set (FVS)

FVS control uses only the topological structure of a network and knowledge of target phenotype biomarkers to induce a phenotype change [30,44]. In FVS control, by manipulating the internal state of the feedback vertex set (i.e. the nodes that intersect every cycle in the network), we disrupt all feedbacks, making the resulting network admit a single steady state, which can be aligned with one of the original system’s dynamic attractors. Thus, a *FVS* of a graph is a minimal set of nodes whose removal leaves the graph without cycles. FVS control has been successfully applied to a variety of networks and has been shown to provide an upper bound on the cardinality of the single set of control nodes needed to reach all attractors [29, 31]. The FVS method’s advantages include: (i) control simply requires fixing the internal state of the FVS to match that of the desired attractor, and (ii) making robust predictions that depend only on the network structure and not on dynamical details. For a transcription factor network underlying a phenotypic switch, the FVS is a set of transcription factors that, when controlled to match the expression of a desired phenotype, will shift the cell towards that phenotype [2].

We formally define a *feedback vertex set* of a directed graph *W* as a possibly empty subset *I* of vertices such that the di-graph *W \ I* is acyclic, where *W \ I* denotes the resulting di-graph when all vertices of *I* are removed from *W*, along with all edges from or towards those vertices. An alternative way to view FVS is as trees and forests. Recall that a *tree* is an undirected graph in which any two vertices are connected by exactly one path, that is, a connected acyclic undirected graph. A *forest* is defined as an undirected graph in which any two vertices are connected by at most one path, that is, an acyclic undirected graph, or a disjoint union of trees [47]. Define a graph *G* = (*V, E*) that consists of a finite set of vertices *V* (*G*) and a set of edges *E*(*G*). Then a FVS of *G* is a subset of vertices *V*′ ⊆ *V* (*G*) such that the removal of *V*′ from *G*, along with all edges incident to *V*′, results in a forest [48]. As such, a FVS must contain all source nodes and a node in every cycle. In other words, a FVS is a set of “determining nodes” such that if the dynamics of the determining nodes are given for large times, then the dynamics of the whole system are determined uniquely for large times [2, 30, 49].

### 3.4 Stable Motifs (SM)

Stable motif (SM) control is based on the identification of self-sustaining generalized positive feedback loops in the dynamic model. Each of these stable motifs determines a region of the state space from which dynamical trajectories cannot escape, called a *trap space*. Further, a stable motif (or a succession of multiple stable motifs) determines a dynamical attractor (i.e. phenotype). There is a SM control set associated with each attractor of the system, and the impact of numerous regulators on a single node can be addressed and analyzed with the method [32]. By definition, a *stable motif* is a strongly connected subgraph of the expanded graph that [2]:

1. contains either a node or its complement but not both
2. contains all inputs of its composite nodes (if any exist)

First, implement the expanded network that is used to add information about the combinatorial interaction and signs of nodes. Composite nodes represent the AND interaction and complementary nodes represent the NOT interaction. Each original node *i* is denoted by *x*_*i*_ in the expanded graph, and a complementary node (∼ *x*_*i*_) is added if the original node represented suppression. Then, all NOT functions are replaced by its appropriate complementary node in the function. Next, edges are included where each edge is a positive regulation, contrary to the original wiring diagram [2, 50].

The second step is to make distinctions between OR rules and AND rules by using composite nodes for functions ivolving ANDs. To do this, the functions must be in disjunctive normal form in order to uniquely determine edges. A special node is included for AND rules, and edges are drawn from the non-composite nodes of the network that form the actual composite rule. It is noted that the benefit of such an action is that the reader is able to see all regulatory functions simply from the topology of the expanded network. Now that the expanded graph is complete, using the definition above we can search for SMs within the network. The group of nodes included in the SM represent partial fixed points, from which the remaining nodes can be calculated using the original Boolean functions [2, 50].

## 4 Efficiency management

In the age of “Big Data”, models are increasingly large and ever more complex. Currently the human genome is estimated to have approximately 25,000 genes, and single genes can encode multiple proteins. What’s more, post-translational modifications add even more complexity to the proteome, with an estimated list of greater than one million proteins [51]. Even networks of merely 100 nodes present a state space much larger than the total estimated cells in the human body [2]. Therefore, the question of control efficiency is an open problem to address. Below, we present possible options for addressing network sizes that are too large for target discovery to be performed in a timely fashion.

### 4.1 Reduction techniques

The magnitude of the BN state space for *n* genes is 2^*n*^. Thus, an increase of GRN size will exponentially increase the computational burden for its analysis, which means brute-force methods for small systems are not sufficient. Many reduction techniques allows to reduce the size of the network while preserving dynamical features (e.g., fixed points and periodic attractors), see [36, 37]. Reduction techniques were implemented in a pancreatic cancer model that effectively decreased the total network size from sixty-nine nodes to twenty-two nodes, a 68% reduction [33]. Critically, when a node was deleted, its function values were substituted directly into its downstream signal recipient(s) to maintain key network communications. Further, nodes containing self-loops cannot be removed, this includes input (source) nodes and self-modulating nodes.

First, nodes with one input and one output were removed, but maintain nodes with self-loops and phenotypes as biomarkers (see Figure 4) [35]. Next, remove nodes with either one input and multiple outputs, or vice versa (see Figure 5). Lastly, remove nodes with low connectivity relative to the remaining nodes (see Figure 6). These techniques have been shown to preserve fixed points but not complex attractors, yet, there are results indicating a conservation of attractors [33, 36, 37]. For an example of one input and one output, consider FGFR from the pancreatic cancer model [33]. The original model’s neighborhood about FGFR is shown in Figure 4a with equations (5) - (6).

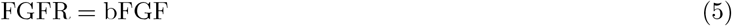

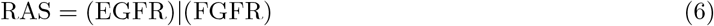

**Figure 4:**
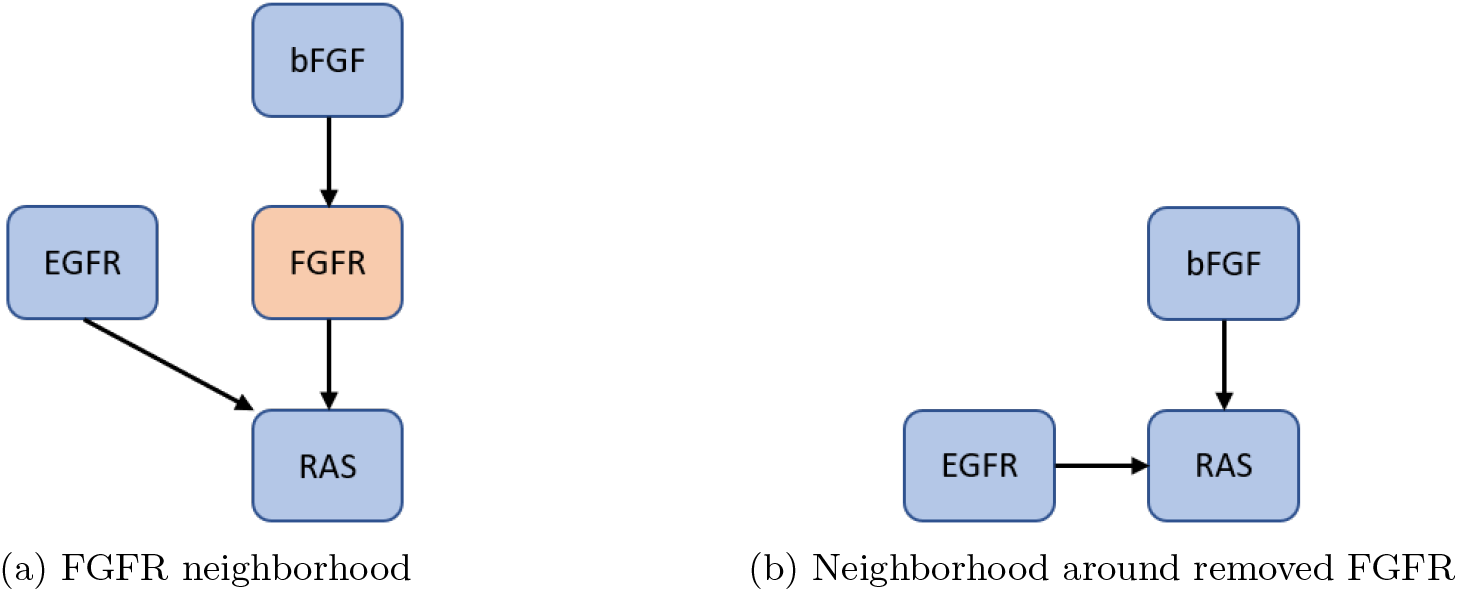
Single-in-single-out removal. Here, we show how to remove FGFR from the network and still maintain downstream signaling. See equations 5 - 8 for functional maintenance.

**Figure 5:**
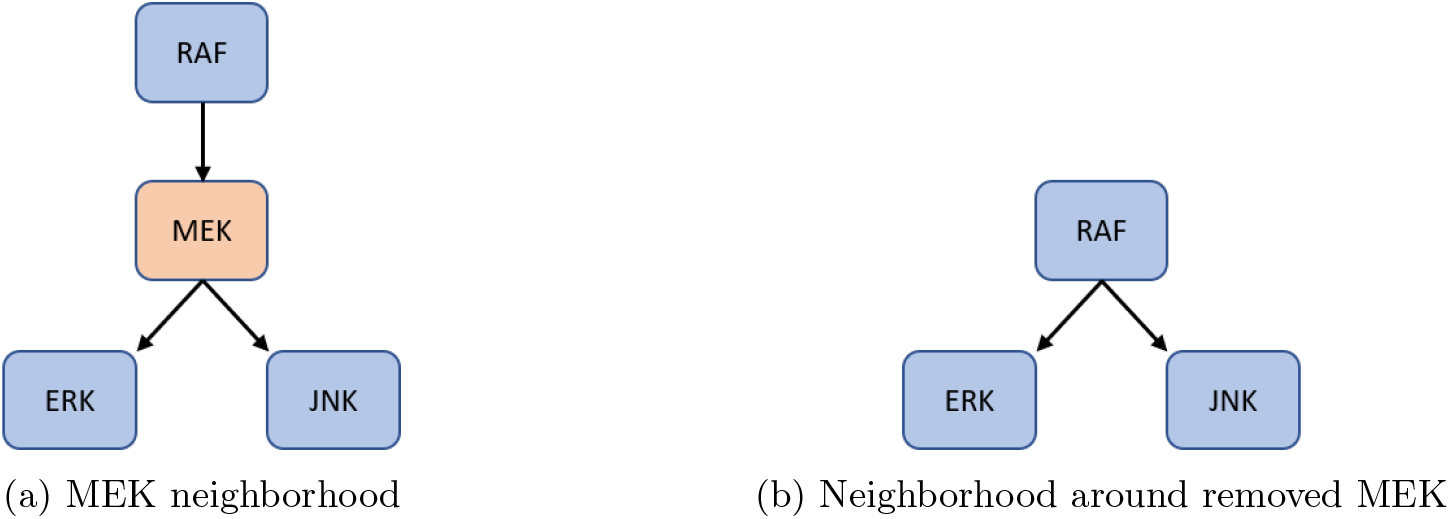
Single-in-multi-out removal. Here, we show how to remove MEK from the network and still maintain downstream signaling. See equations 9 - 14 for functional maintenance.

**Figure 6:**
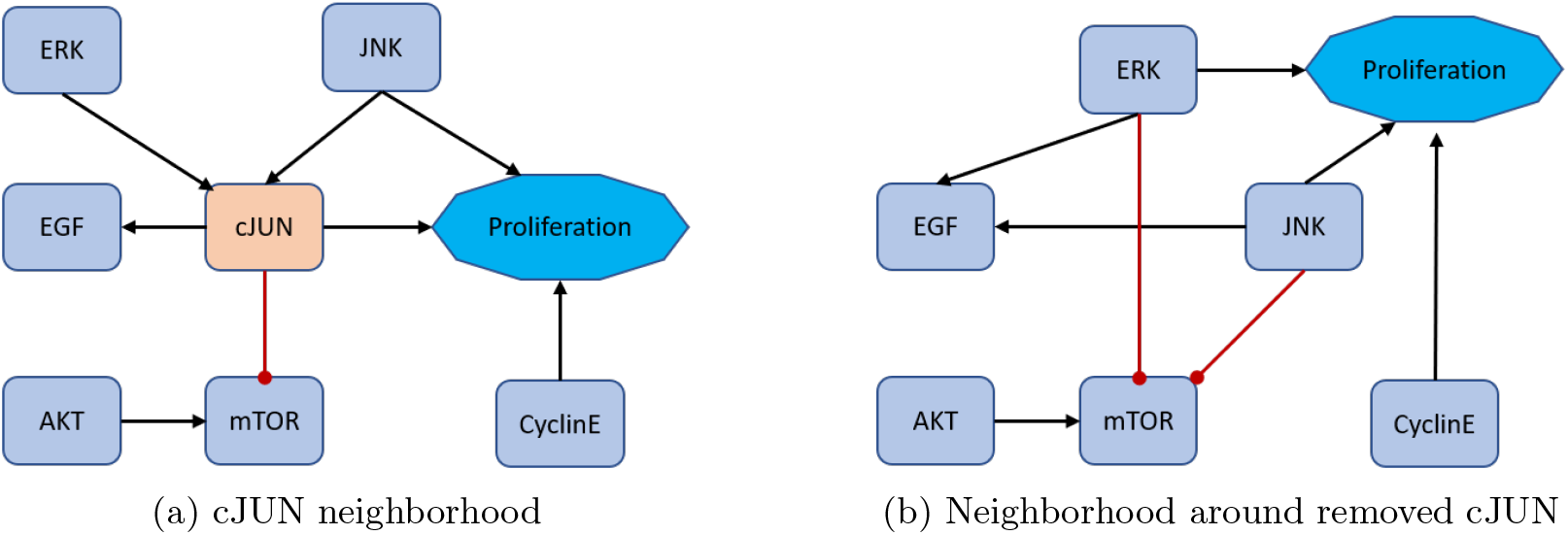
Low connectivity removal. Here, we show how to remove cJUN from the network and still maintain downstream signaling. See equations 15 - 22 for functional maintenance.

After reduction, we obtain the neighborhood seen in Figure 4b with equations (7) - (8).

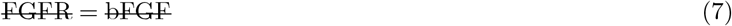

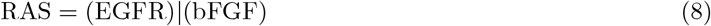

For an example of either one input and multiple outputs, or vice versa, consider MEK from [33]. The original model’s neighborhood about MEK is shown in Figure 5a with equations (9) - (11).

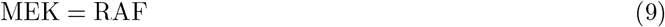

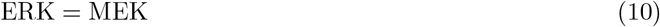

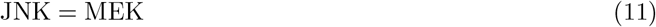

After reduction, we obtain the neighborhood seen in Figure 5b with equations (12) - (14).

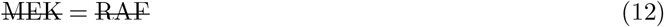

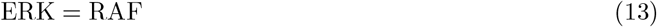

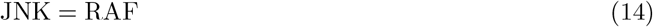

Lastly, for an example low connectivity removal, consider cJUN [33]. The original model’s neighborhood about cJUN is shown in Figure 6a with equations (15) - (18).

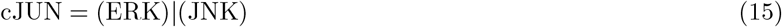

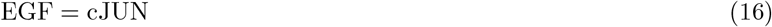

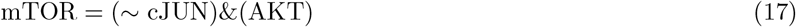

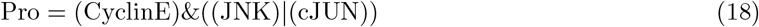

After reduction, we obtain the neighborhood seen in Figure 6b with equations (19) - (22).

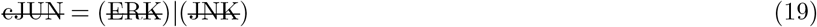

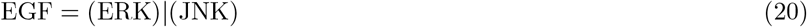

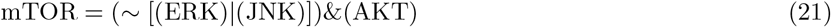

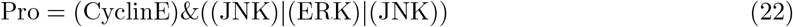

### 4.2 Modularity techniques

Systems biology is capable of building complicated structures from simpler building blocks, even though these simple blocks (i.e. modules) traditionally are not clearly defined. The concept of modularity detailed in [34] is structural by nature, in that, a *module* of a BN is a subnetwork in which the restriction of the network to the variables of a subgraph has a strongly connected wiring diagram. This framework introduces both a structural and dynamic decomposition that encapsulates the dynamics of the whole system simply from the dynamics of its modules. Consequently, the decomposition yields a hierarchy among modules that can be used to specify controls. That is, by controlling key modules we are able to control the entire network [2].

Within the modularity framework, the dynamics of the state-space for Boolean network *F* are denoted as 𝒟(*F*), which is a collection of all minimal subsets of attractors, *A*, satisfying *F* (*A*) = *A*. Further, if *F* is decomposable (say into subnetworks *H* and *G*), then we can write *F* = *H* × *G* which is called the *coupling* of *H* and *G*. In the case where the dynamics of *G* are dependent on *H*, we call *G non-autonomous*, denoted as 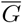. Then we adopt the following notation: let *A* = *A*_1_ ⊕ *A*_2_ be a set of attractors of *F* with *A*_1_ ∈ 𝒟(*H*) and 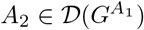 [2].

For an example, consider the network in Figure 7a with

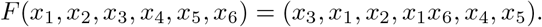

**Figure 7:**
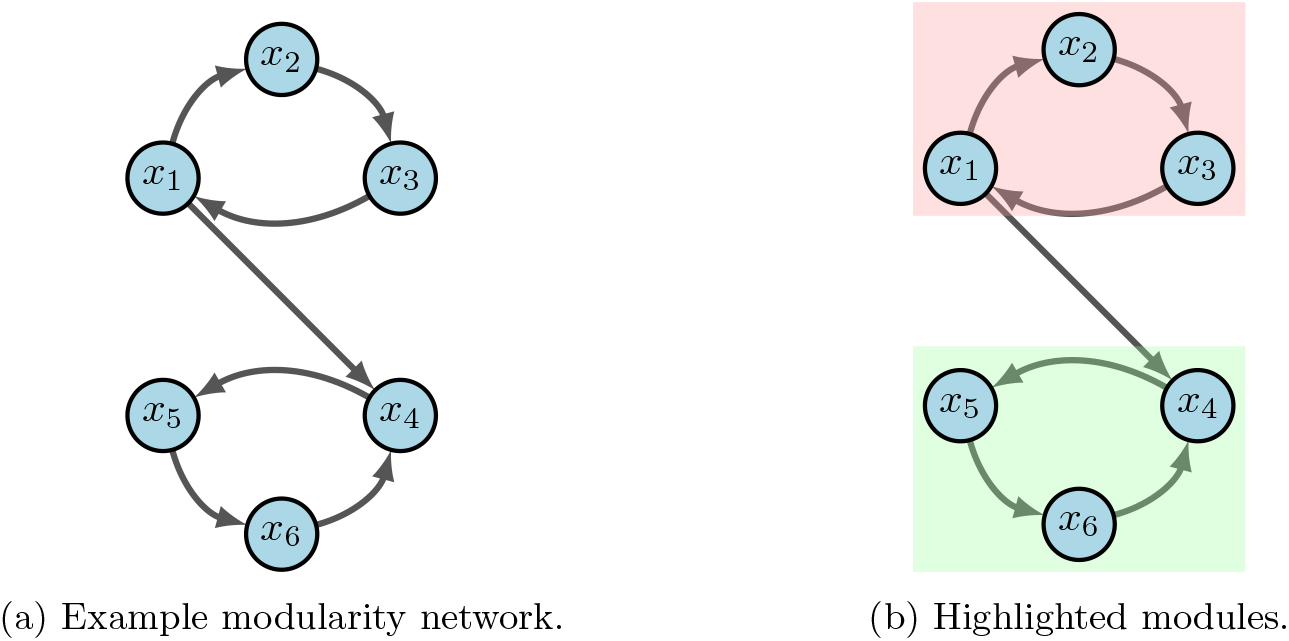
Modularity example [2].

From the given wiring diagram, we derive two SCCs where module one (red in 7b) flows into module two (green in 7b). That is, *F* = *F*_1_ × *F*_2_ with

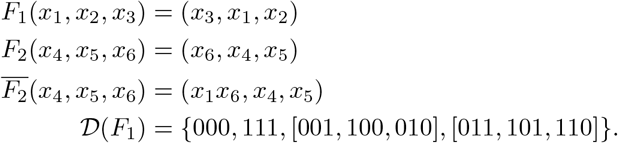

Suppose we aim to stabilize the system into *y* = 000000. First we see that either *x*_1_ = 0, *x*_2_ = 0 or *x*_3_ = 0 stabilize module one (i.e. *F*_1_) to *A*_1_ = 000 by applying the FVS method from Section 3.3. Likewise, *x*_4_ = 0, *x*_5_ = 0 or *x*_6_ = 0 stabilize module two (i.e. 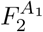) to *A*_2_ = 000. Thus, we conclude that *u* = (*x*_1_ = 0, *x*_6_ = 0) achieves the desired result [2].

## 5 Limitations

Even though phenotype control theory shows massive potential, the field overall has some limitations, along with those of each technique we have described. From a biological and translational perspective, it remains yet to be validated as a viable option for clinical application. Further, the human genome is highly complex, with signaling mechanisms that are far from well understood. This leads modelers to rely on speculative networks and hypothesized functional communication rules.

Regardless of method, each of the resulting outputs are merely theoretical controls and must be parsed to find tangible targets (or combinations of targets). Efficacy of the resulting targets can be established computationally, which is discussed in the Appendix 7.3. The parsing process can include brute-force testing of all controls, knowledge of the regulatory network topology, knowledge of literature pertaining to particular controls, or a mixture of various techniques [22, 26]. Some controls may not be biologically achievable, others may be insufficient if applied independently, while some simply do not perform as desired.

Since we do not apply optimal control, another constraint to address is how to select controls that prioritize certain interventions over others. These criteria might include selection according to effectiveness (e.g. shorter absorption time), total/side effects (e.g. number of changes in the original state space), target “depth” within the network, and practical implementability. Many of the selection criteria will need stochasticity (such as for time to absorption), which can be achieved via SDDS [10, 52] or asynchronous simulations (see Appendix 7.2).

When it comes to network reduction, techniques can prove extremely tedious if networks are notably large. Further, the reduction techniques can change the long-term outlooks of key analytical features such as cyclical attractors. It has been shown that the methods in Section 4.1 will maintain fixed points, but they do not necessarily maintain cyclical attractors [33, 36, 37]. Even though examples have been shown to maintain all attractors [2, 33], one can easily show counter examples that do not (see the small T-LGL model in Section 2.1). Thus, a fully developed methodology for efficient reduction is yet to be seen, which could be important for analyzing large models.

Additionally, computational complexity varies across methods. For instance, the CA method makes use of computing Gr*ö*bner bases for a system of polynomials and, depending on the algorithm used, it has been shown to have doubly exponential complexity [26]. However, GRNs with small sets of regulatory nodes can compute Gr*ö*bner bases in a reasonable time [26, 53].

For CK, the problem of finding the minimal set of controlling nodes was shown to be NP-hard [54], and the problem of the existence of multiple possible minimal control sets is NP-complete. Thus, when computing CKs, no algorithm is expected to run faster in the worst case than checking every possible subset of increasing size, since the rounds of pinning to find CK’s are representative of NP-hard problems. Moreover, the average CK sizes scale logarithmically with the number of attractors [29].

The computational time to find a single FVS is reasonable, the issue arises when trying to find all possible FVSs. The global stabilization of BNs have been shown to have computational complexity that is exponential with respect to the number of state variables [55, 56]. However, while the problem of exactly identifying the minimal FVS has complexity of NP-hard, a variety of fast algorithms exist to find close-to-minimal solutions [31, 57].

Lastly, the complexity of calculating SMs using the domain of influence (DOI), through the expanded graph [50], is bounded by the order of the sum the number of nodes and edges in the expanded network, *O*(*N*_*ex*_ +*E*_*ex*_). Subsequent calculations for finding control sets from the DOI become more complex. So called “well behaved degree distribution” networks give calculated order *O*(*k*^2^*N* ^2^), where *k* are the regulators for each node *N*. Those networks considered to have “skewed degree distribution” are bounded by *O*(*N*^3^) [50].

## 6 Conclusions

In this paper, we reviewed various techniques for implementing target discovery and control of gene regulatory networks. Due to the growing nature of the field, there are always emerging, novel techniques to implement and we acknowledge that the methods included here are not fully exhaustive [58–62]. Even so, we have set out to provide a list of varying options, depending on the specific aims and information available to users, that represent a broad range of applicable theory. We also hope to spark conversations and ideas for solving open problems in the field, as well as inspire application of these concepts across a wide range of disciplines, not strictly biology.

In addition to toy examples for each method (see Appendix), we also applied each approach to a well known cancer model (T-LGL Leukemia) to explore overlaps and differences among the processes. In particular, we showed that FVS provides an upper bound for the amount of targets needed to achieve network control, whereas CA and CK can provide minimal sets. Perhaps the most versatile method shown is CA, where users have wide ranging options to personalize their search (i.e. nodes vs. edges, use existing attractors, generate new attractors, and block transitions or regions). These overlaps have also been shown in a computational pancreatic cancer model [22, 33].

Even though there is not a common theoretical framework to apply all methods, we do see that each is capable of affirming discoveries across other methods while also suggesting possible novel targets of their own. We believe the future is bright for synthetic modeling and control of cell signaling networks, and the methods reviewed here in are just the beginning.

## Acknowledgements

The authors would like to thank Reinhard Laubenbacher and Reka Albert for their discussions and suggestions during in the initial stage of this project. Further, DP was supported by the NIH Training Grant T32CA165990. D.M. was partially supported by a Collaboration grant (850896) from the Simons Foundation.

## 7 Appendix

### 7.1 Elementary examples for control methods

#### 7.1.1 Computational Algebra

Consider the network in Figure 8, with the following regulatory functions.

**Figure 8:**
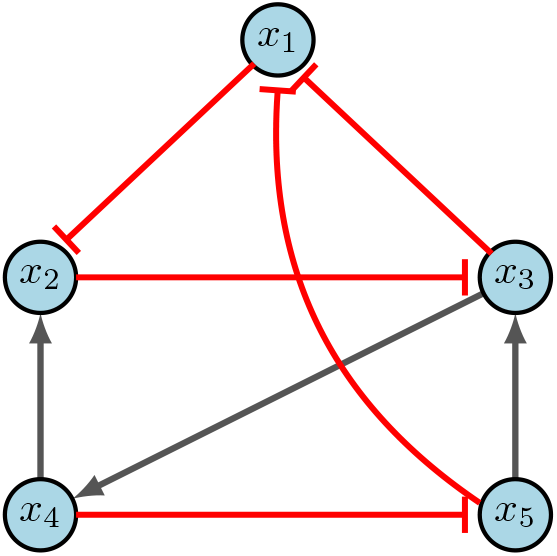
CA example [2].

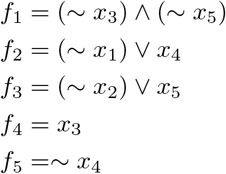

Using Table 5, we rewrite our functions as the following simplified polynomials.

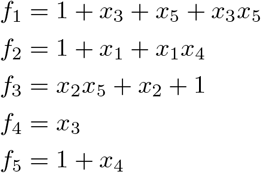

We can then find the fixed points of the system by solving *f*_*i*_ = *x*_*i*_ for *i* = 1, …, 5. Another way to view this step is as finding roots of *g*_*i*_ = 0 where *g*_*i*_ = *f*_*i*_ − *x*_*i*_, then finding the Grobner basis of the ideal *I* = ⟨*g*_1_, …, *g*_5_⟩. In any case, the example in Figure 8 does not contain any fixed points. However, further state space analysis does reveal two attractors: {01011, 01100} and {00101, 01010, 01110, 01111, 10001, 11000}. Now, we encode our edge controls as

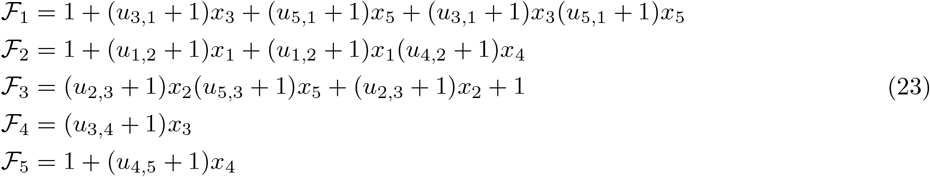

and node controls as

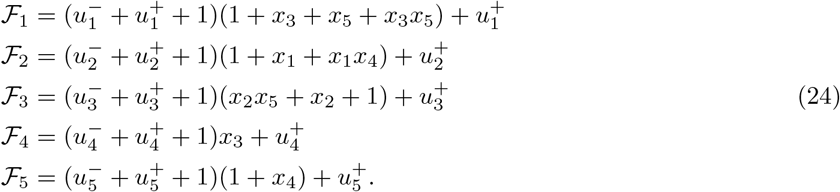

Let’s consider the objective of generating new attractors, and assume we want our steady state to be *y* = 11110. In general, one can search the entire system for controls, but there may be special cases where limiting decisions can be made amongst collaborators. For arguments sake, suppose we want to find edge knockouts and limit our search to edges *x*_3_ → *x*_1_, *x*_5_ → *x*_1_, and *x*_2_ → *x*_3_. Then the updated edge equations (Eq. 23) become

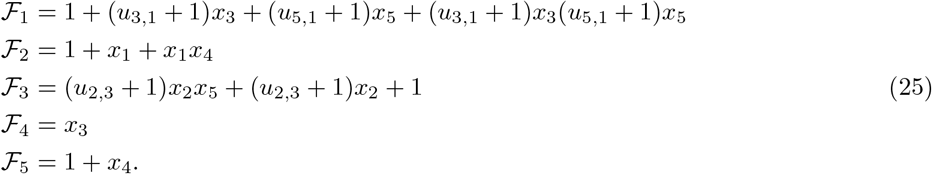

Evaluating at *y* = 11110 yields

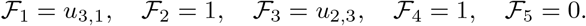

Therefore, the desired fixed point is achieved if and only if *u*_3,1_ = *u*_2,3_ = 1. That is, the controls for *u*_3,1_ and *u*_2,3_ are *active*, such that we must delete both corresponding edges. Similarly, we can determine node control to achieve new fixed point *y* = 11110. Again, for simplicity, we limit ourselves to *x*_1_ knock-in, *x*_3_ knock-out and knock-in, and *x*_4_ knock-in. The updated node equations (Eq. 24) then become

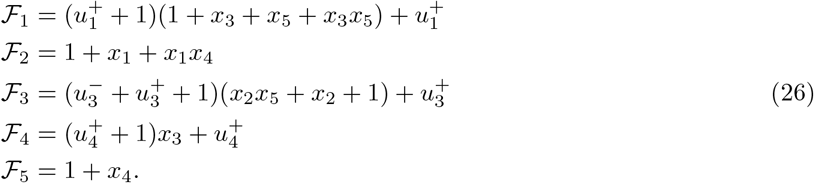

Evaluating at *y* = 11110 yields

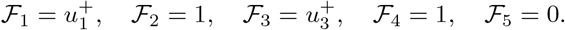

Thus, the desired fixed point is achieved if and only if 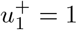 and 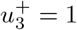. Importantly, this means that the controls by themselves are insufficient but together they achieve the desired goal. One can easily see that requiring numerous controls in much larger systems may not be biological feasible, which is why alternate objectives can prove useful.

Suppose we determine that *y* = 01111 is in a diseased attractor which we want to destroy. We can then aim to block the transition from *y* to *F* (*y*) = 01110. We limit ourselves to considering edges from *x*_3_ → *x*_1_, *x*_5_ → *x*_1_, *x*_3_ → *x*_4_, and *x*_4_ → *x*_5_. The updated edge equations (Eq. 23) become

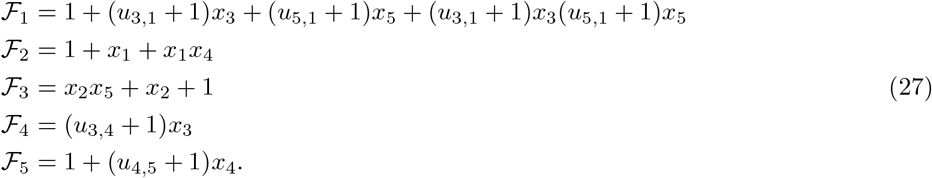

Evaluating at *y* = 01111 yields

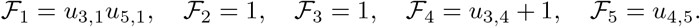

This means that Eq. 2 becomes

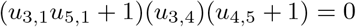

giving three possible solutions: *u*_3,1_ = *u*_5,1_ = 1, *u*_3,4_ = 0, or *u*_4,5_ = 1. Notice that we again have a combinatorial solution in *u*_3,1_, *u*_5,1_ since they are insufficient individually but successful together, *u*_3,4_ = 0 means that the control is inactive, and *u*_4,5_ is a singleton control.

Lastly, consider the objective of region blocking. Suppose we want to avoid regions where *x*_3_ = 0, and we will limit ourselves to nodes *x*_2_ knock-out, *x*_3_ knock-in, and *x*_4_ knock-in. Then the updated node equations (Eq. 24) become

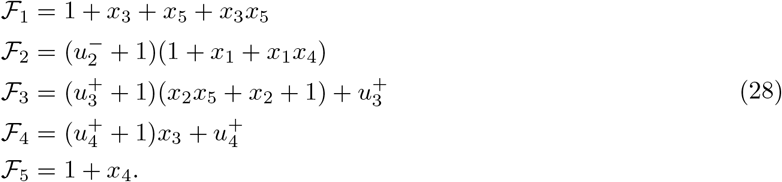

Next, we see that Eq. 3 yields

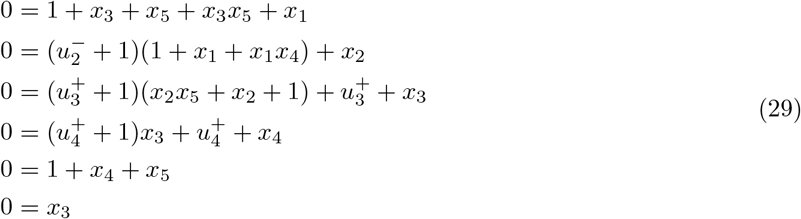

Using computation algebra tools to compute the Grobner basis of the ideal associated to the above equations, we encode the system of equations to achieve the ideal:

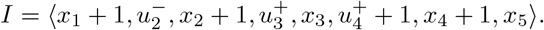

This means the original system has the same solutions as the following system.

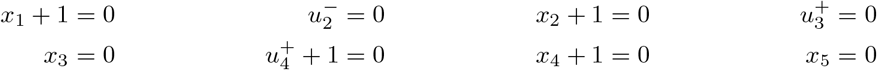

Recall that our goal is to block the region *x*_3_ = 0 by finding parameters that guarantee the above system has no solutions. Utilizing equations that only contain control parameters we have 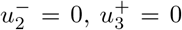, and 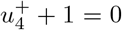. Thus, if we allow either 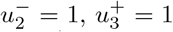, or 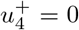, then our system will have no solution, as needed. Since *x*_3_ is limiting criteria and 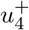 is an inactive control, that leaves 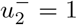 as the desired target. As one can see, the computational algebra method is quite versatile [2].

#### 7.1.2 Control Kernel

Consider the network in Figure 9. Steady state analysis reveals two fixed points: 000100 and 111011. Suppose our control objective is *x*_4_ = 0, which is the second fixed point respectively. We first notice that there are no input nodes, which means we move on to distinguishing nodes. Then the CK method (correctly) indicates that *x*_1_ = 1 will direct the system into the desired fixed point. Admittedly, while the CK method is straight forward, the software used to implement the search can be difficult to navigate [2].

**Figure 9:**
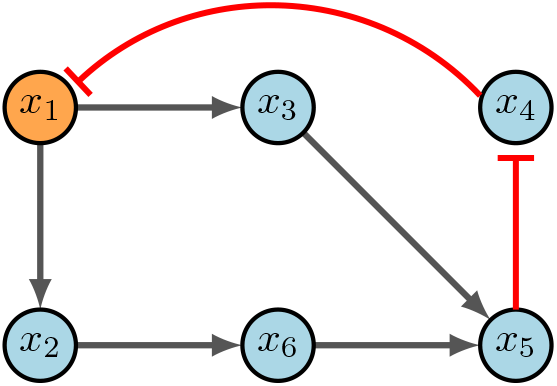
CK example [2].

#### 7.1.3 Feedback Vertex Set

Figure 10 contains a simple example of identifying a FVS. The input node (*x*_1_) is always in the control set, while the only other node required is one of those in the 3-cycle. As scene in the figure, 10a is the example wiring diagram and 10b - 10d show the three possible FVS’s. One can easily see that the strategy for FVS is quite simple, yet, it can produce larger control sets than necessary. Further, we may not obtain all FVS’s if the system has many attractors [2].

**Figure 10:**
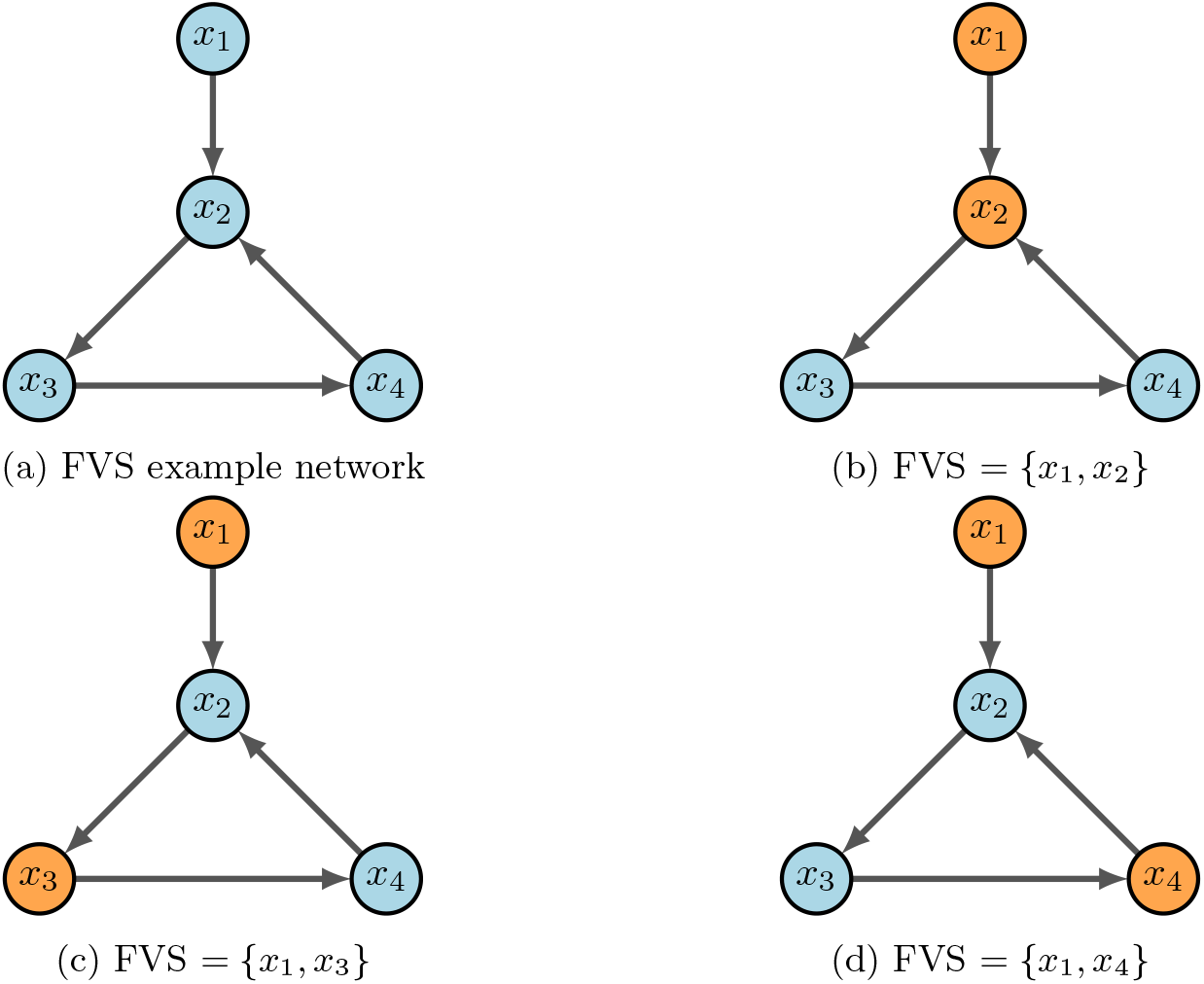
FVS example [2].

#### 7.1.4 Stable Motifs

Consider the example network in Figure 11a, with the following functions and negated functions.

**Figure 11:**
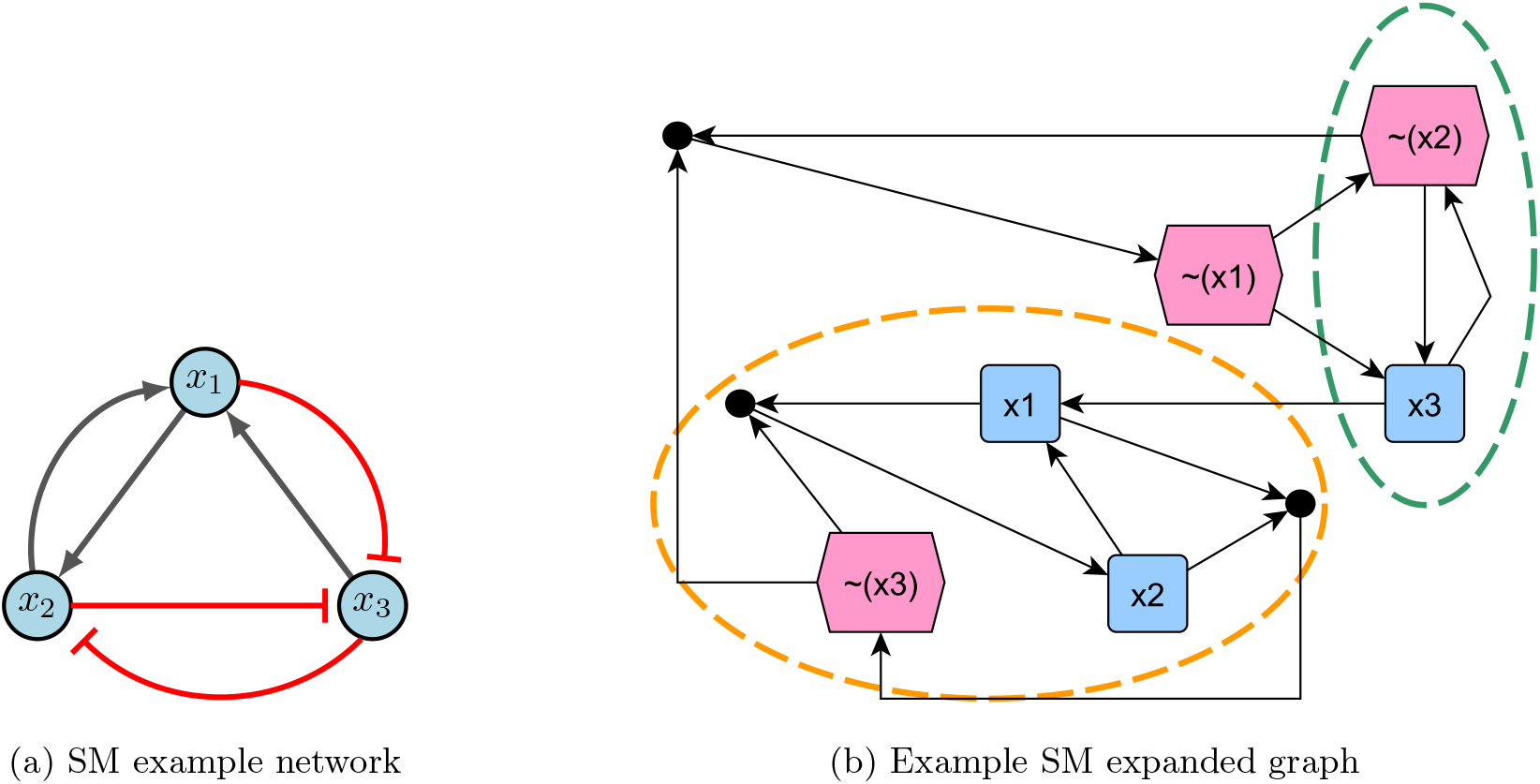
Stable motif example [2].

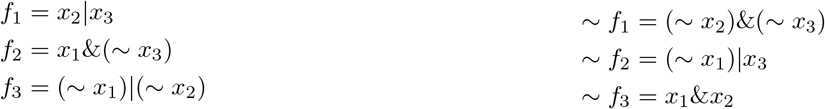

Using the aforementioned steps, the expanded graph obtained is Figure 11b. Notice there are two stable motifs (circled in orange and green), which indicate a fixed point (110) and a partial fixed point (*X*01). To find the rest of partial fixed point, substitute known values into the original functions. Therefore,

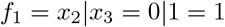

which gives 101 as the second fixed point. Since the control sets are subsets of the stable motifs, we have {*x*_2_ = 1, *x*_3_ = 0} or {*x*_1_ = 1, *x*_3_ = 0} for fixed point 110, and {*x*_2_ = 0} or {*x*_3_ = 1} for fixed point 101 [2].

### 7.2 Finite Dynamical Systems

For the last few decades, a popular modeling approach for gene regulation has been to implement dynamical systems over finite fields. Here, functions can be interpreted as modeling information processing within cells, which determines cellular behavior. As depicted in Figure 12, 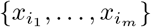 represent the input genes or predictor genes, 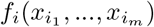 is the internal update function or predictor rule, and *x*_*i*_ is the target gene.

**Figure 12:**
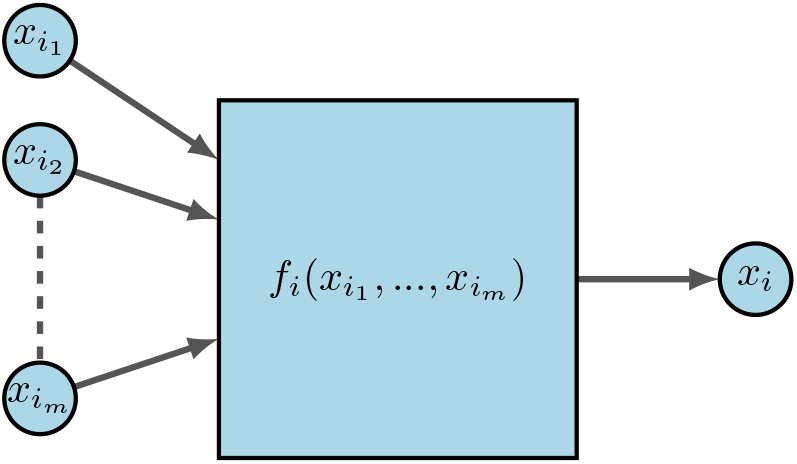
FDS for gene regulation [2].

First, let *X* = *X*_1_ × *X*_2_ ⋯ × *X*_*n*_ be the Cartesian product of finite sets. A *local model* over a finite set *X* is an *n*-tuple of coordinate functions *F* = (*f*_1_, *f*_2_ …, *f*_*n*_), where *f*_*i*_ : *X*^*n*^ → *X*. Each function *f*_*i*_ uniquely determines a function

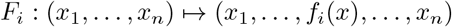

and *x* = (*x*_1_, …, *x*_*n*_). Every local model defines a canonical *finite dynamical system (FDS)* map, where the functions are updated as

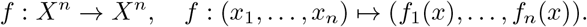

Note that discrete does not necessarily imply finite. Take the natural numbers ℕ = 1, 2, 3, 4, …, for example. The set is clearly discrete, yet its cardinality is infinite. In general, we cannot always write a function as a tuple if the space is simply “discrete”. In order to provide structure to each *X*_*i*_, we embed *X*_*i*_ into a finite field where, for some prime *p*,

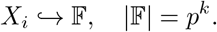

For example, if we desire states of Low, Medium, and High to represent levels of gene expression, then *X*_*i*_ = {*L, M, H*} ⤷ *𝔽*_3_ = {0, 1, 2}. We call these *mixed-state* models when states are non-binary. For the case when states are binary (i.e. ON or OFF, HIGH or LOW, 1 or 0), we call these models *Boolean networks* [2].

#### 7.2.1 Boolean Networks

Boolean networks (BNs) are popular because we can build effective models without the use of constants or rates. This then eliminates the need for tedious parameter discovery. Rather, BNs focus on the mechanics and logic of the system. BN models were originally introduced in 1963 by Kauffman and Thomas to provide a coarse grained description of gene regulatory networks [23, 63]. Within a BN there are three main components: structure (wiring diagram), functions (regulatory rules), and dynamics (attractors). As we begin to define our terms, it may be helpful to keep Figure 13 in mind as a basic example. Given *n* binary variables, define a *Boolean Network* as an *n*-tuple of coordinate functions

**Figure 13:**
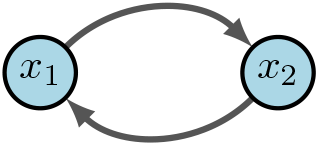
Simple Boolean network [2].

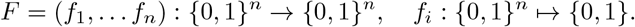

The *wiring diagram* of *F*, call it *W*, is then defined as a directed graph with *n* nodes {*x*_1_, *x*_2_, …, *x*_*n*_} such that there is an edge in *W* from *x*_*j*_ to *x*_*i*_ if *f*_*i*_ depends on *x*_*j*_. That is,

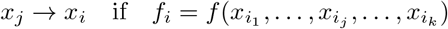

Within *W* we denote positive edges as *x*_*j*_ → *x*_*i*_ and negative edges as *x*_*j*_ ⊣ *x*_*i*_ (or sometimes *x*_*j*_ ⊸ *x*_*i*_). Biologically, a positive edge is representative of activation while a negative edge represents inhibition. For example, in Figure 13 we see the wiring diagram of *F* = (*f*_1_, *f*_2_) = (*x*_2_, *x*_1_).

Now that we have structure and functions, the dynamics of *F* are traditionally described as: (1) trajectories for all 2^*n*^ possible initial conditions, or (2) a directed graph with nodes in 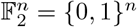. In the first case, a *trajectory* is a sequence 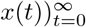 given by the difference equations *x*(*t* + 1) = *F* (*x*(*t*)) for all *t* ≥ 0 [34]. For example, Figure 13 would yield deterministic trajectories

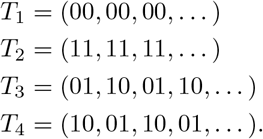

The *phase space* (also called state space) of *F* is the directed graph with vertex set *S*^*n*^ and edge set {(*s, f* (*s*))|*s* ∈ *S*^*n*^}. Simply put, in a BN, *S* is the set of all possible states, and their respective transitions according to the model *F* form the state space (see Figure 14). A node *s* ∈ *S* is called *transient* if *f*^*k*^(*s*) ≠ *s* for all *k >* 1, a node *s* ∈ *S* is called *periodic* (or cyclic) if *f*^*k*^(*s*) = *s* for some *k* ≥ 1, and a node *s* ∈ *S* is called a *fixed point* if *f* (*s*) = *s*. We can also think of the phase space as having strongly connected components (SCCs), where a SCC is said to be *terminal* if it has no out-going edges. Thus, a transient state is not in a terminal SCC, a cyclic attractor is in a terminal *k*-cycle (*k* = 1 is a fixed point), and any instance of an SCC otherwise is a *complex attractor*. In other words, we define an *attractor* as a set of states from which there is no escape as the system evolves, and an attractor with a single state is called a fixed point. Thus, given sufficient time, the dynamics of a BN always end up in a fixed point or (complex) attractor.

**Figure 14:**
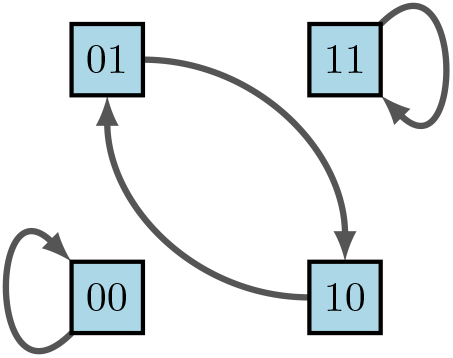
Phase space of diagram 13 [2].

For example, it was previously shown above that *F* = (*f*_1_, *f*_2_) = (*x*_2_, *x*_1_). To find the dynamics of the corresponding state space *S* = {00, 01, 10, 11}, one can construct truth Table 4 using lexicographic ordering. It is important to point out that we denote the states in order of the variable so that

**Table 4:** Dynamic truth table for Figure 13.

| $x_1$ | $x_2$ | $f_1 = x_2$ | $f_2 = x_1$ |
| --- | --- | --- | --- |
| 0 | 0 | 0 | 0 |
| 0 | 1 | 1 | 0 |
| 1 | 0 | 0 | 1 |
| 1 | 1 | 1 | 1 |

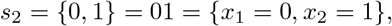

because maintaining order is highly important for correct interpretation of state values. The left columns indicate the possible states of our nodes *x*_1_ and *x*_2_, whereas the right columns indicate their deterministic updates according to the functions *f*_1_ and *f*_2_. Therefore, from the framework we see in Figure 14 that we have two fixed points and one cycle.

Up to this point we have only discussed linear BNs, but real-world models are almost always highly nonlinear (see Figure 15). To accommodate these nonlinear regulatory networks, we implement various classes of functions based on three main Boolean logical rules - AND, OR, NOT. Some use XOR (exclusive OR), but for simplicity it is excluded here. Assume the variables *x* and *y* are given in a BN. Then Table 5 summarizes the functionality and notation used for each of the three main rules.

**Figure 15:**
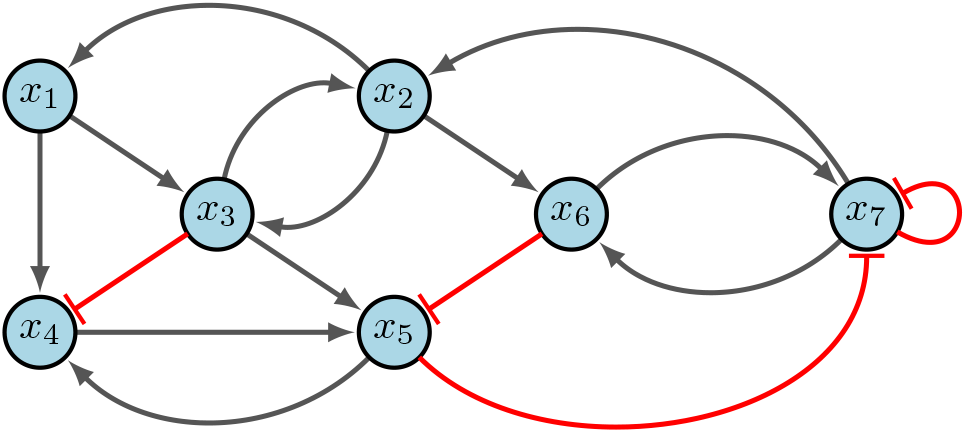
Nonlinear Boolean network [2].

**Table 5:**
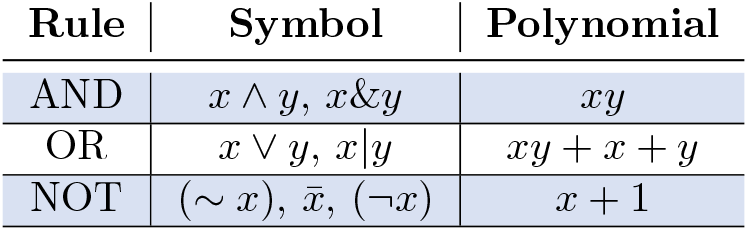
Standard Boolean logical rules.

A common criticism of using discrete models for regulatory networks such as BNs is that deterministic dynamics are artificial. In reality biological systems do not contain a “central clock”, but instead the concentration levels of gene products change and respond to stimuli on varying time-scales. Thus, the update schedules chosen play a significant role in the accuracy of the model. *Synchronous* update schedules produce deterministic dynamics, wherein nodes are all updated simultaneously so that

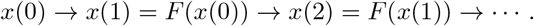

On the other hand, *asynchronous* update schedules produce stochastic dynamics, wherein a randomly selected node is updated at each time step so that

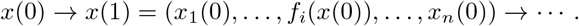

Lastly, *sequential* update schedules are performed asynchronously according to a designated permutation *σ* = (*σ*_1_, …, *σ*_*n*_) of (1, …, *n*). Specifically, if we define *F*_*i*_(*x*_1_, …, *x*_*n*_) = (*x*_1_, …, *f*_*i*_(*x*), …, *x*_*n*_), then the update is given by

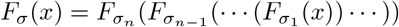

according to the order designated by *σ*. This is sometimes done when the ordering of gene updates are known, as some may update faster than others. For example, using our simple example in Figure 13, Figure 16 shows the varying impacts of these three update schedules.

**Figure 16:**
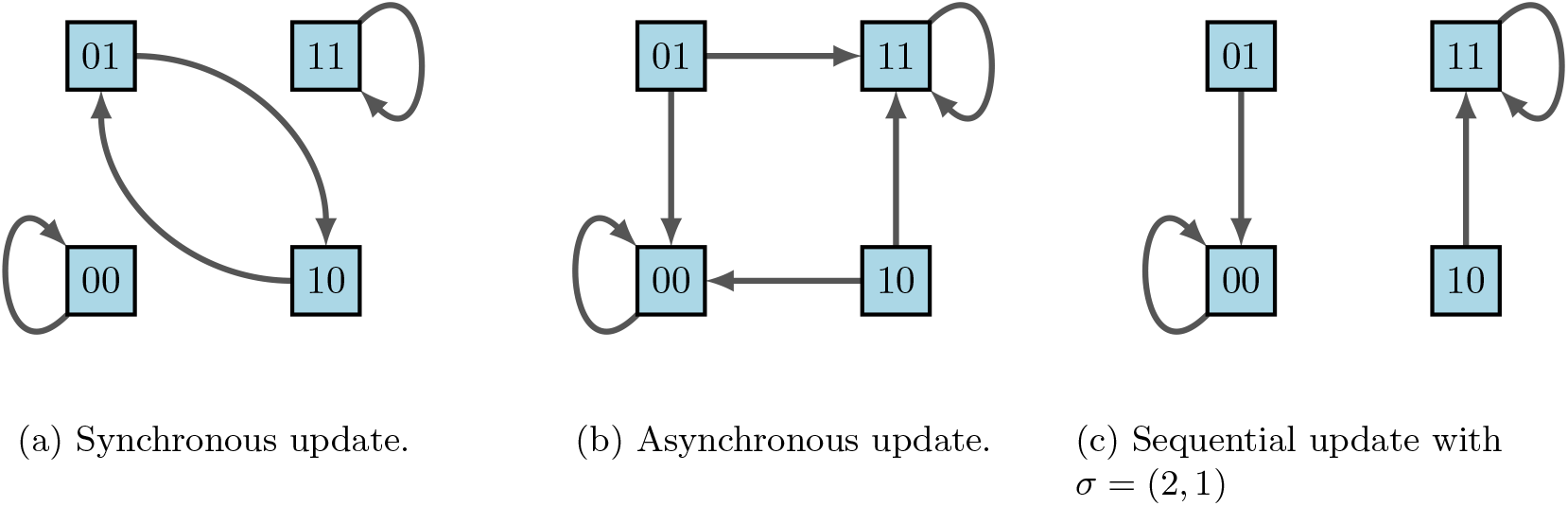
State-space dynamical variants according to update schedules [2].

We can easily observe from Figure 16 that fixed points are maintained across all update schedules. However, cycles are not necessarily preserved. This is where the framework of Stochastic Discrete Dynamical Systems (SDDS) is beneficial [2,11,22,33]. Developed in [11], SDDS incorporates Markov chain tools to study long-term dynamics of Boolean networks. SDDS uses parameters based on designated propensities to model node (and pathway) signal activation and deactivation, also referred to as degradation. In essence, SDDS merges the synchronous and asynchronous update schedules described above. One propensity is used when the update positively impacts the node, in the sense that the node increases its value from OFF to ON. Another propensity is used when the update negatively affects the node in the sense that the node decreases its value from ON to OFF. More precisely, an SDDS of the variables (*x*_1_, *x*_2_, …, *x*_*n*_) is a collection of *n* triples

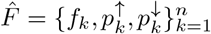

where for *k* = 1, …, *n*,

- *f*_*k*_ : {0, 1}^*n*^ → {0, 1} is the update function for *x*_*k*_
- 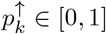 is the activation propensity
- 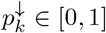 is the deactivation propensity

Here, the parameters 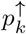 and 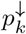 introduce stochasticity. For example, an activation of *x*_*k*_(*t*) at the next time step (i.e. *x*_*k*_(*t*) = 0, *f*_*k*_(*x*_1_(*t*), …, *x*_*n*_(*t*)) = 1, and *x*_*k*_(*t* + 1) = 1) occurs with probability 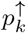. An SDDS can be represented as a Markov Chain via its transition matrix, which can be viewed as transition probabilities between various states of the network. Elements of the transition matrix *A* are determined as follows: consider the set *S* = {0, 1}^*n*^ consisting of all possible states of the network. Suppose *x* = (*x*_1_, …, *x*_*n*_) ∈ *S* and *y* = (*y*_1_, …, *y*_*n*_) ∈ *S*. Then, the probability of transitioning from *x* to *y* is

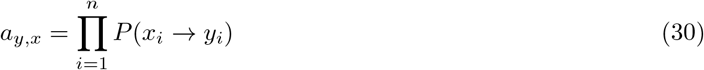

where entries are stored column-wise and

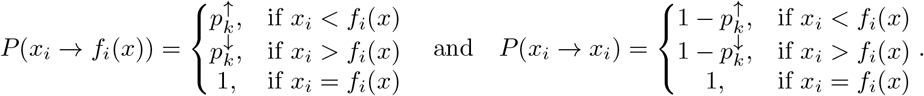

It follows that *P*(*x*_*i*_ → *y*_*i*_) = 0 for any *y*_*i*_ ∉ {*x*_*i*_, *f*_*i*_(*x*)}. Therefore, we achieve *A* = [*a*_*y,x*_]_*x,y*∈*S*_. Note that when propensities are set to *p* = 1, we have a traditional BN. With this framework, we built a simulator that takes random initial states as inputs and then tracks the trajectory of each node through time. Long-term phenotype expression probabilities can then be estimated, as well as network dynamics with (and without) controls [2].

### 7.3 Simulating Target Efficacy

To determine the efficacy of controls, we compare uncontrolled simulations with the appropriate target control simulations. Thus, a good control will produce low disease levels and high health levels [2]. We can do so by utilizing a stochastic simulator based on SDDS [2,11,22,33], which requires several inputs before it can begin. The number of input variables in each Boolean function is given by the vector *nv*. Next, we need the variables for each gene in the form of an *m* × *n* matrix called *varF* where *m* is the maximum number of inputs, *n* is the number of genes, and information is stored column-wise. The number of variables will vary between functions. Since only the first *nv*(*i*) elements of the *i*th column are relevant, all remaining entries are set as (−1). Now we construct the truth table *F* in compact form with size 2^*m*^ × *n*. Again, the length of each column *i* will vary but only the first 2^*nv*(*i*)^ entries are relevant. So all remaining entries are set as (−1). It is vitally important to maintain numerical ordering, which is why the columns of *F* are in lexicographic binary arrays [25].

We must also establish propensities in the form of a 2 × *n* matrix *c* that contains values for 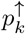 and 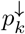. The values chosen for propensities may perturb results, as we saw in Figure 16. But for all intents and purposes, we typically use 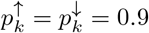 (i.e. follow the function rules 90% of the time). Finally, we can run simulations using inputs: *F, varF, nv, number of states (usually Boolean), c, n, number of steps*, and *number of random initializations*. We have also implemented versions that allow for mutation induction and specified initial states. As a result, we achieve time-course trajectories, and we can use the Markov chain structure of SDDS to analyze features such as time to absorption, stationary distributions, and more.

As an example, consider the simple 3-cycle in Figure 17. This particular system has two fixed points ({000} and {111}) as well as two attractors ({001, 100, 010} and {011, 101, 110}). Simulations were conducted using the variables in Table 6, with 1000 random initializations, 100 time steps (function updates), and injecting 1% noise. The overall state-space is shown in Figure 18. In Figure 19a, the uncontrolled simulation shows the oscillatory nature of attractors. However, Figures 19b and 19c show that inducing control on *x*_1_ is enough to drive the system to one fixed point or the other. Therefore, the SDDS simulator has the ability to show long-term trajectories and impact of controls over time.

**Table 6:**
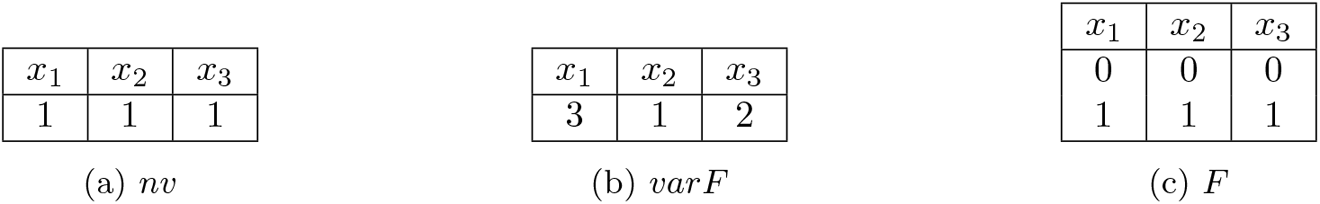
Variable tables for simple 3-cycle simulations in Figure 17 [2].

**Figure 17:**
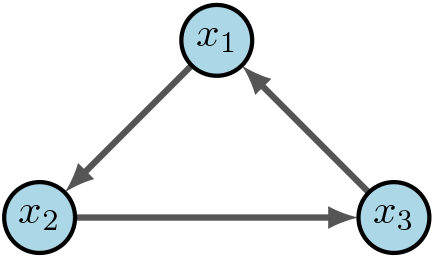
Simple 3-cycle [2].

**Figure 18:**
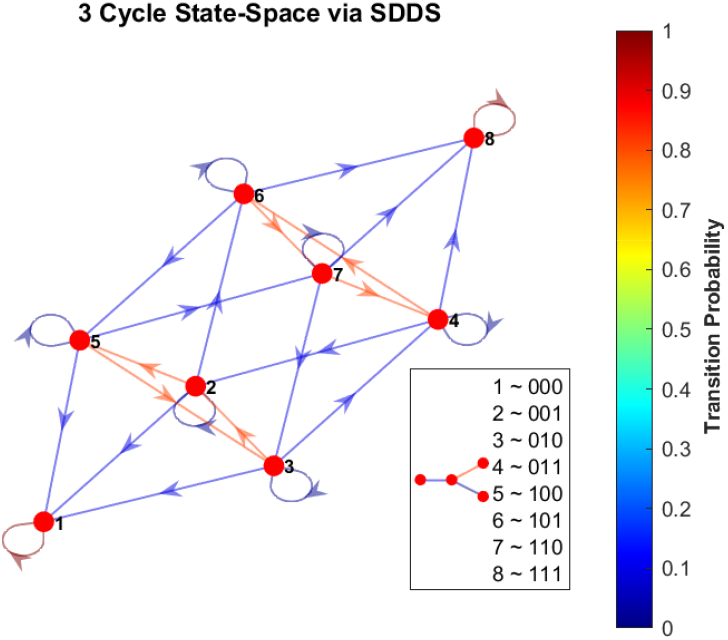
Phase-space of simple 3-cycle. Here we show the state-space of the example from Figure 17, using SDDS with transition probabilities, with nodes written in lexicographical ordering.

**Figure 19:**
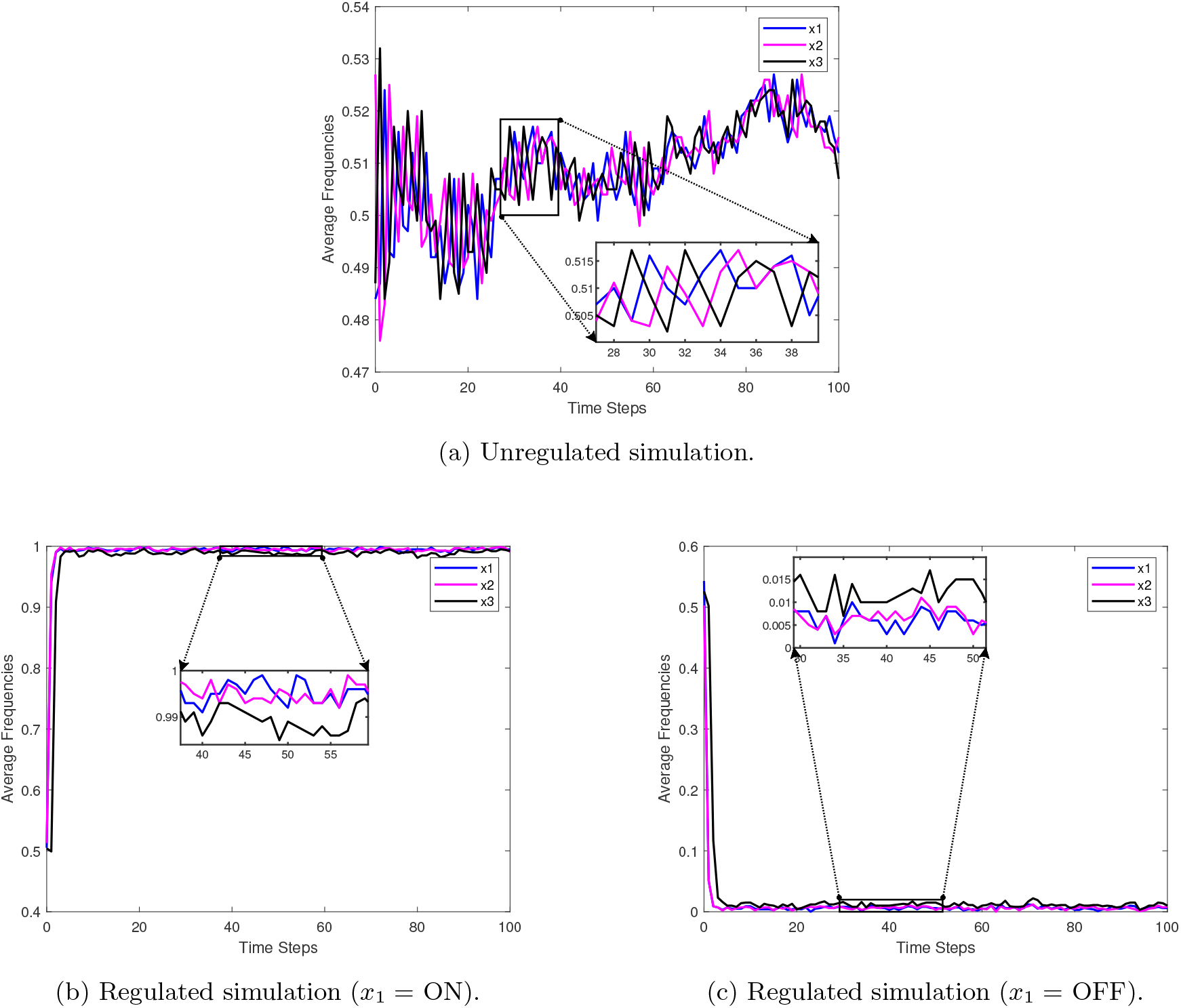
Simulation examples for a simple 3-cycle with 1% noise [2].

### 7.4 Software

- Cumulative files for all control techniques and examples, as well as “how-to” documentation [2]
  – https://github.com/drplaugher/SMATA_pipeline
- CA: used to find fixed points, controls, and run simulations [22, 33, 64]
  – use the example files above
  – see also, https://github.com/drplaugher/PCC_Mutations
- CK: used to find control kernels [29]
  – https://doi.org/10.5281/zenodo.5172898
- FVS: used to find FVSs [30, 44]
  – https://github.com/jgtz/FVS_python3
- Modularity: used to find strongly connected components (modules) [34]
  – use the example files above
- SM: used to find stable motifs and dynamic attractors [32, 65]
  – https://github.com/jgtz/StableMotifs
  – https://github.com/jcrozum/pystablemotifs

### 7.5 Supplementary Tables

**Table 7:** Small T-LGL rules

| Node | Boolean Rule |
| --- | --- |
| S1P | (not Ceramide) |
| FLIP | (not DISC) |
| Fas | (not S1P) |
| Ceramide | Fas and (not S1P) |
| DISC | Ceramide or (Fas and (not FLIP)) |
| Apoptosis | DISC |

**Table 8:**
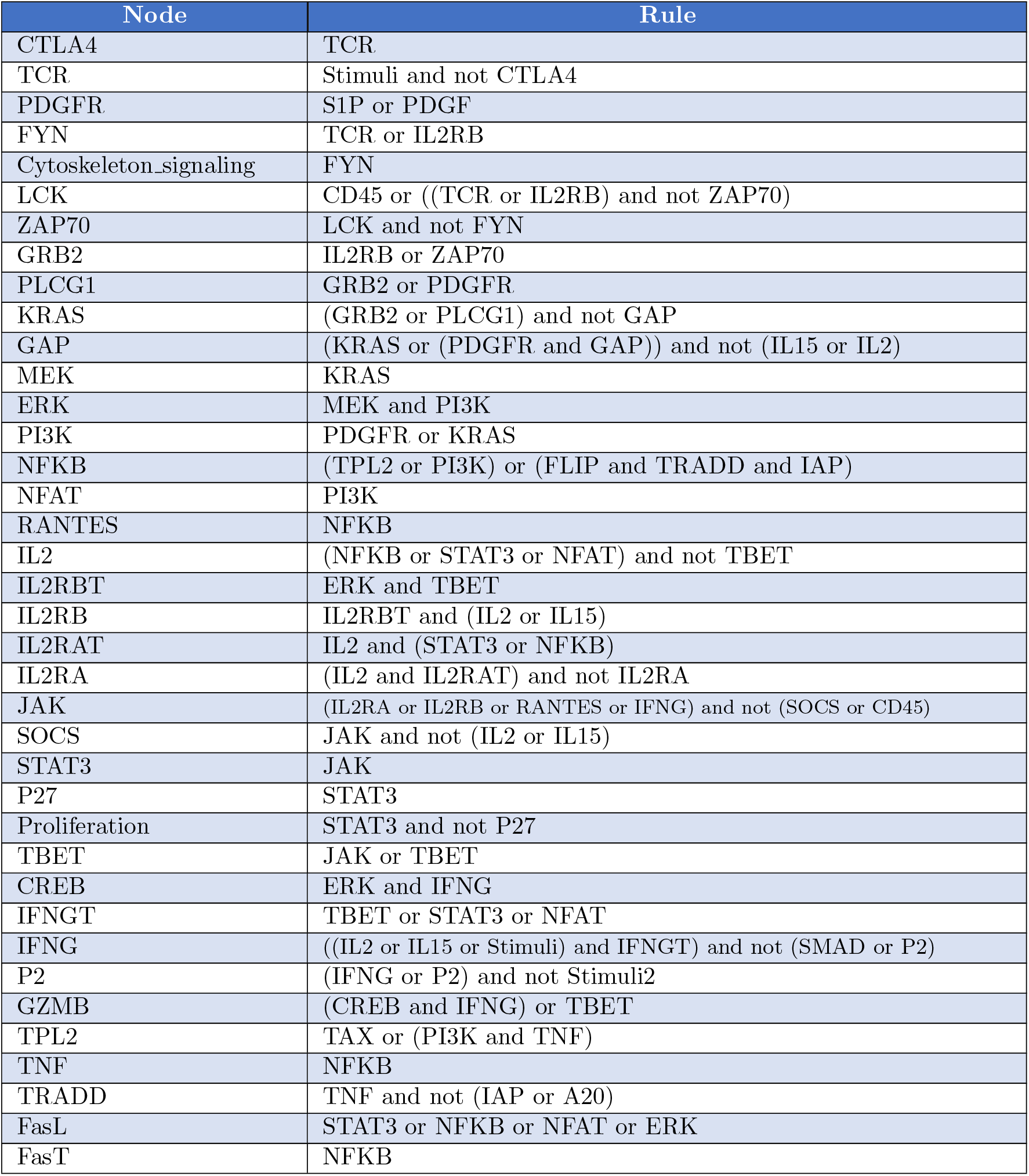
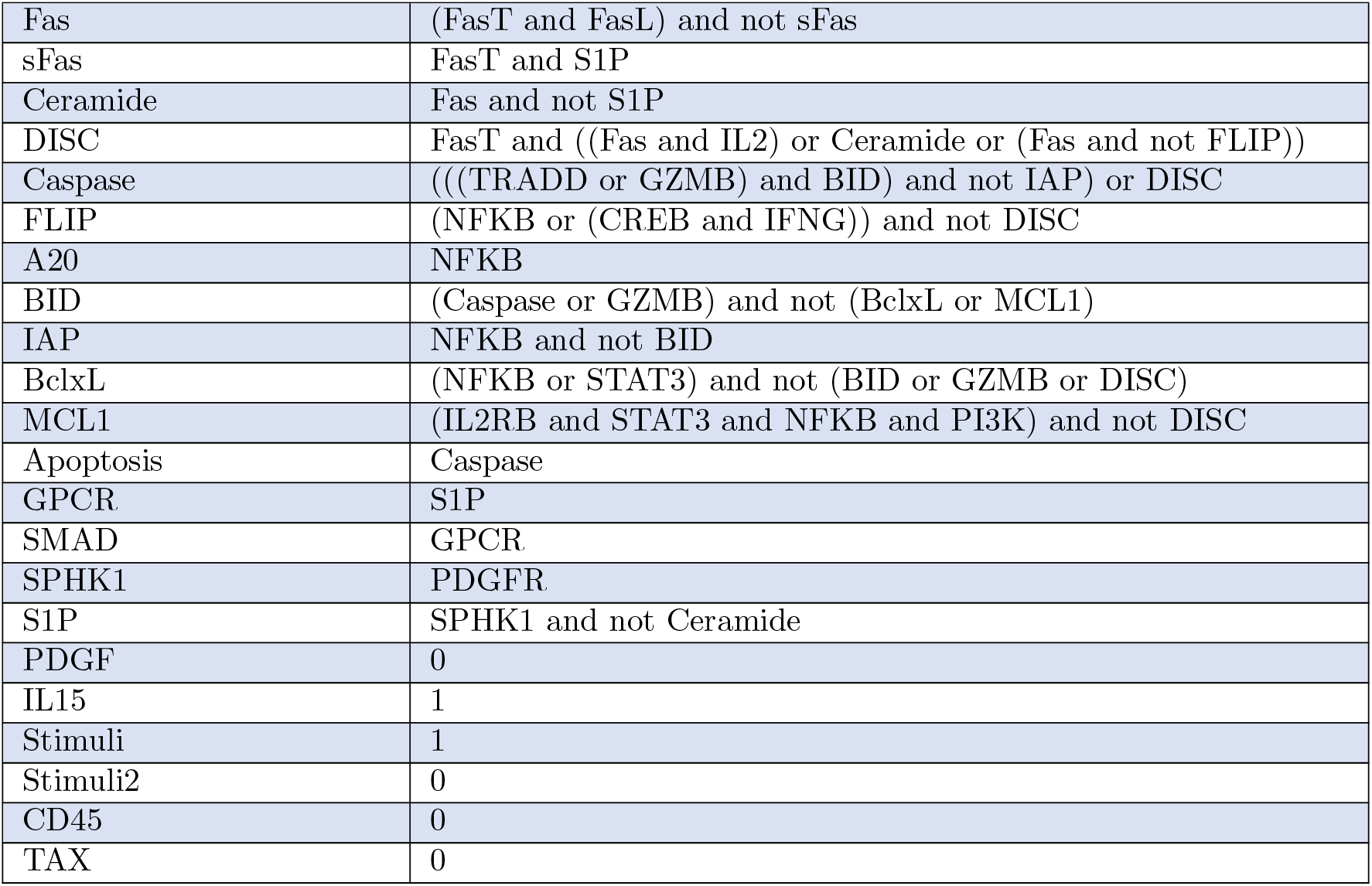
Functions for large T-LGL model.

## Notes

### Competing Interest Statement

The authors have declared no competing interest.

